# Molecular and spatial analysis of ganglion cells on retinal flatmounts: diversity, topography, and perivascularity

**DOI:** 10.1101/2024.12.15.628587

**Authors:** Nicole Y. Tsai, Kushal Nimkar, Mengya Zhao, Matthew R. Lum, Yujuan Yi, Tavita R. Garrett, Yixiao Wang, Kenichi Toma, Franklin Caval-Holme, Nikhil Reddy, Aliza T. Ehrlich, Arnold R. Kriegstein, Michael Tri H. Do, Benjamin Sivyer, Karthik Shekhar, Xin Duan

## Abstract

Diverse retinal ganglion cells (RGCs) transmit distinct visual features from the eye to the brain. Recent studies have categorized RGCs into 45 types in mice based on transcriptomic profiles, showing strong alignment with morphological and electrophysiological properties. However, little is known about how these types are spatially arranged on the two-dimensional retinal surface—an organization that influences visual encoding—and how their local microenvironments impact development and neurodegenerative responses. To address this gap, we optimized a workflow combining imaging-based spatial transcriptomics (MERFISH) and immunohistochemical co-staining on thin flatmount retinal sections. We used computational methods to register *en face* somata distributions of all molecularly defined RGC types. More than 75% (34/45) of types exhibited non-uniform distributions, likely reflecting adaptations of the retina’s anatomy to the animal’s visual environment. By analyzing the local neighborhoods of each cell, we identified perivascular RGCs located near blood vessels. Seven RGC types are enriched in the perivascular niche, including members of intrinsically photosensitive RGC (ipRGC) and direction-selective RGC (DSGC) subclasses. Orthologous human RGC counterparts of perivascular types - Melanopsin-enriched ipRGCs and ON DSGCs - were also proximal to blood vessels, suggesting their perivascularity may be evolutionarily conserved. Following optic nerve crush in mice, the perivascular M1-ipRGCs and ON DSGCs showed preferential survival, suggesting that proximity to blood vessels may render cell-extrinsic neuroprotection to RGCs through an mTOR-independent mechanism. Overall, our work offers a resource characterizing the spatial profiles of RGC types, enabling future studies of retinal development, physiology, and neurodegeneration at individual neuron type resolution across the two-dimensional space.

## INTRODUCTION

Cataloging neuronal types, defining their spatial organization, and mapping neuron-neuron connectivity are essential steps for understanding the assembly and function of neural circuits (Luo et al., 2018; Zeng and Sanes, 2017). The mammalian retina is an ideal model system for these studies due to its compact structure, experimental accessibility, and complex neural network comprised of multiple neuronal types (Kerschensteiner and Feller, 2024; Masland, 2001). Over decades of research, the mouse retina has been found to contain ∼130 distinct neuronal types (Sanes and Masland, 2015; Shekhar and Sanes, 2021), organized across three somatic layers (Masland, 2012).

The innermost layer, known as the ganglion cell layer (GCL), houses the somata of retinal ganglion cells (RGCs), which are glutamatergic projection neurons. RGCs are the retina’s sole output neurons, and their axons communicate visual information to the rest of the brain via the optic nerve (**Figures 1A, B**). Investigating the neurobiology of retinal ganglion cells (RGCs) is crucial for advancing approaches to restore vision in cases of neurological injuries or ophthalmological diseases.

**Figure 1.**
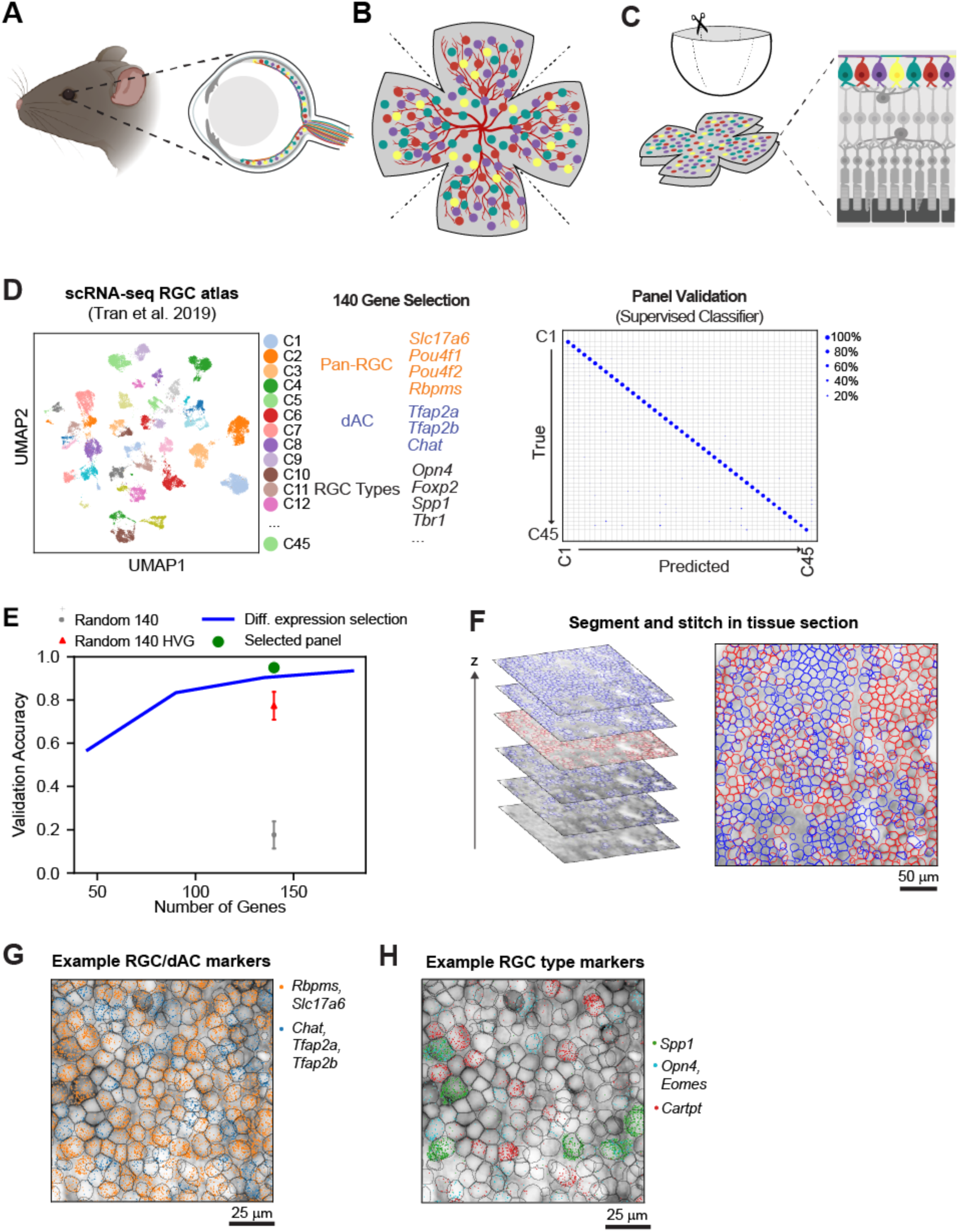
Spatial transcriptomics of the ganglion cell layer (GCL) in thin retinal flatmount sections. A. Illustration of mouse retina showing retinal ganglion cell (RGC) types (colors) in the GCL. Axons of RGCs project through the optic nerve to the rest of the brain. B. Schematic of a GCL flatmount showing RGCs and displaced amacrine cells (dACs) in various colors and non-neuronal cells, including blood vessels (NNs, red). C. Schematic of flatmount preparation. The dissected retina is flattened on filter paper via relieving cuts and cryo-sectioned into ∼12μm thick sections. RGC types are highlighted in flatmount and sectional views. D. UMAP visualization of adult RGC diversity based on scRNA-seq atlas (Tran et al., 2019), MERFISH gene panel consisting of 140 genes optimized for RGC classification based on scRNA-seq atlas, and confusion matrix showing the performance of an XGBoost classifier trained on these genes. E. Validation accuracies of XGBoost classifier trained on various gene selection approaches. The validation of our 140-gene panel was compared to a 140-gene panel from random subsets of 2000 highly variable genes (HVGs). In addition, varying numbers of genes (*45, 90, 135, 180*) were selected by picking differentially expressed genes from each of the 45 clusters. Error bars: ±1 SD (n=10 random gene subsets). F. Overview of segmentation workflow. Cellpose is applied to identify cell somata from optical sections (left), which are stitched together for the final segmentation (right). Red denotes cells identified in the third optical section, while blue denotes additional cells identified in the other optical sections. G. Transcripts are visualized in an individual tissue section with the segmentation masks. Dots correspond to unique transcripts. Transcript colors correspond to marker genes for RGCs (*Rbpms, Slc17a6;* orange) and dACs (*Chat, Tfap2a, Tfap2b;* blue). H. As in G, highlighting transcripts for specific groups of RGC types. α-RGCs (*Spp1*), ipRGCs (*Opn4* and *Eomes*), ON-OFF DSGCs (*Cartpt*).

The RGC class comprises diverse neuronal types, most of which respond selectively to specific visual features, such as motion, orientation, or edges, channeling visual signals into multiple parallel information streams (Gollisch and Meister, 2010). Recent studies have classified ∼45 RGC types in mice based on molecular profiles and demonstrated their alignment with dendritic morphologies and visual responses (Baden et al., 2016; Goetz et al., 2022; Huang et al., 2022; Rheaume et al., 2018; Tran et al., 2019). Advances in mouse transgenics (Badea et al., 2009; Duan et al., 2014; Huberman et al., 2008; Kim et al., 2010; Siegert et al., 2009) and single-cell RNA-sequencing (scRNA-seq) (Rheaume et al., 2018; Tran et al., 2019) have furnished markers for selectively labeling and manipulating specific RGC types. Experiments leveraging these molecular or genetic markers have enabled the mapping of synaptic connectivity within the retina and onto retinorecipient regions (Duan et al., 2014; Krishnaswamy et al., 2015; Martersteck et al., 2017; Tsai et al., 2022). Crucially, these markers have served as a basis for unifying classification schemes (Goetz et al., 2022; Huang et al., 2022; Tran et al., 2019). Furthermore, the ability to distinguish RGC types based on their molecular profiles has also been helpful in studying their development (Shekhar et al., 2022; Whitney et al., 2023), evolutionary conservation (Hahn et al., 2023), and type-specific differences in responses to optic nerve injury and neurodegenerative insults (Bray et al., 2019; Duan et al., 2015; Jacobi et al., 2022; Li et al., 2022; Tran et al., 2019; Zhao et al., 2023).

Although RGC classification is nearly complete in mice and quite advanced in several other species (Hahn et al., 2023), questions persist regarding the spatial distribution of different RGC types across the retinal surface and potential variations in their local microenvironments within the retina.

Additionally, for most RGC types, it remains unclear whether their inter-somal spacing follows a non-random pattern—a characteristic known as “mosaicism” that has been documented in several retinal cell types, including some RGC types (Wassle and Riemann, 1978). Conventional scRNA-seq techniques discard spatial information due to the requirement of enzymatic dissociation of the tissue.

Recently, the application of serial section electron microscopy on intact blocks of the retina has provided detailed anatomical data for RGC types, including soma distributions and dendritic architecture (Bae et al., 2018; Briggman et al., 2011). However, this technique has limited throughput (∼1mm^2^ per sample), covering only ∼5% of the retinal surface. On the other hand, transgenic markers and immunohistochemical labeling (Bleckert et al., 2014; Heukamp et al., 2020; Rousso et al., 2016) have examined full spatial distributions of select RGC subsets. Still, these techniques are limited by multiplexing constraints and potential variability from transgene positioning effects (Duan et al., 2018). These technical barriers underscore the need for approaches to profile RGC diversity with spatial information over the entire retinal surface.

To map the spatial organization of RGC types across the retina, we developed a protocol integrating spatial transcriptomics with immunohistochemical labeling on flatmount retinal sections. Together with computational analyses, these datasets enabled us to map the spatial distributions of all transcriptomically defined RGC types across the entire retinal surface. Many RGC types exhibit biased distributions of somas along the dorsoventral or nasotemporal axes and exhibit non-random inter-soma spacing indicative of mosaic arrangements. Additionally, we identified seven RGC types that are frequently found in proximity to retinal blood vessels, including members of the intrinsically photosensitive RGC (ipRGC) and direction-selective ganglion cell (DSGC) subclasses. We validated the perivascular properties of these ipRGC and DSGC subtypes using transgenic lines specific to them. In addition, we provided anatomical evidence that this association is conserved in the human retina. Finally, we found that the ipRGC and DSGC perivascular types marked by these transgenics exhibited preferential neuronal survival following optic nerve crush, suggesting a neuroprotective effect associated with perivascularity.

## RESULTS

### Spatial transcriptomic imaging of retinal flatmount sections

The mouse GCL, a ∼20µm thick monolayer, contains the somata of RGCs, displaced amacrine cells (dACs), and non-neuronal cells (NNs) such as astrocytes, endothelial cells, pericytes, and microglia. Endothelial cells, pericytes, and other NNs are assembled into the superficial layer (SL) of the retinal vasculature within the same space, spanning the two-dimensional surface of the retina (**Figures 1A, 1B**). As RGCs form a flat layer of neurons in the GCL, the flatmount preparations provide comprehensive access to them. We developed an integrated experimental and computational workflow to visualize RGC diversity across the entire retina.

Our method is based on multiplexed-error robust fluorescent *in situ* hybridization (MERFISH), an imaging-based spatial transcriptomics approach (Chen et al., 2015; Moffitt and Zhuang, 2016) that visualizes individual transcripts at ∼1 µm resolution (**Methods**). Since MERFISH requires thin tissue sections, we optimized a protocol to obtain a series of 12µm thick flatmount sections to cover the entire GCL *en face* (**Figure 1C**). We performed MERFISH imaging on these sections, targeting 140 genes optimized for classifying RGC types based on their transcriptomic profiles from our established scRNA-seq atlas (Tran et al., 2019) (**Figure 1D, Table S1**). Although the current gene panel covers only a small fraction of the transcriptome, simulations using the scRNA-seq atlas verified that these 140 genes reliably distinguish all 45 RGC types, with ∼94% precision and ∼91% recall (**Figures 1D, 1E, S1A-S1C; Methods**).

A single retina yielded four consecutive sections, which were processed using a standardized MERFISH imaging protocol. Cell somata within each section were segmented using a neural network model based on a cell body stain (Figure 1F, **S1D, Methods**). The neural network model was based on Cellpose 2.0 (Stringer et al., 2021), but customized for our dataset through extensive training on manually curated and annotated images (**Methods**). Detected transcripts were assigned to the segmented somata, providing an estimate of the mRNA composition of each detected cell (Figures 1G, 1H). We then aligned the images from the four sections via rigid rotations and translations to reconstruct the full GCL in two dimensions for each retina (**Figure S1E**). We computed a cell-by-gene expression matrix (GEM) for each retina for downstream clustering analyses and visualization. On average, we captured ∼62,000 ± 9,800 cells per retina, assigning each cell an (*x, y*) spatial coordinate within the retinal surface (**Figures S1F, S1G**).

### Separation of major cell classes in the ganglion cell layer

Initial clustering of the data from all six biological replicates separated cells by their class identity (Figures 2A-2C; Methods). Approximately 93% of captured cells were neurons, while 7% were NNs (Figure 2D). Among neurons, 89% were reliably assigned based on their expression of marker genes to the two major classes known to populate the GCL: 49% were RGCs (28,208 ± 3,682 per retina), and 39% were dACs (22,730 ± 4,814 per retina) (Figures 2D**, S2A**). The relative proportion of RGCs among GCL neurons (∼49%) aligns favorably with prior estimates via axon counting (∼44%) (Jeon et al., 1998). Moreover, these estimates suggest that our method captures approximately 63% of RGCs in the GCL per retina (Jeon et al., 1998). The remaining 12% of neurons formed a single ambiguous cluster of low-quality cells (Figures 2D**, S2B**). These cells were excluded from downstream analyses. To validate our manual assignments, we used a tree-based classifier (Chen and Guestrin, 2016) trained on multiple scRNA-seq atlases representing major retinal cell classes (Benhar et al., 2023; Tran et al., 2019; Yan et al., 2020a) (**Figure S2C, Methods**). The classifier confirmed our assignments of RGCs, dACs, and NNs based on marker gene expression. The classifier also verified the absence of bipolar cells (BCs), photoreceptors (PRs), and horizontal cells (HCs), which are retinal neuronal classes not found in the GCL under normal conditions (Figure 2E).

**Figure 2.**
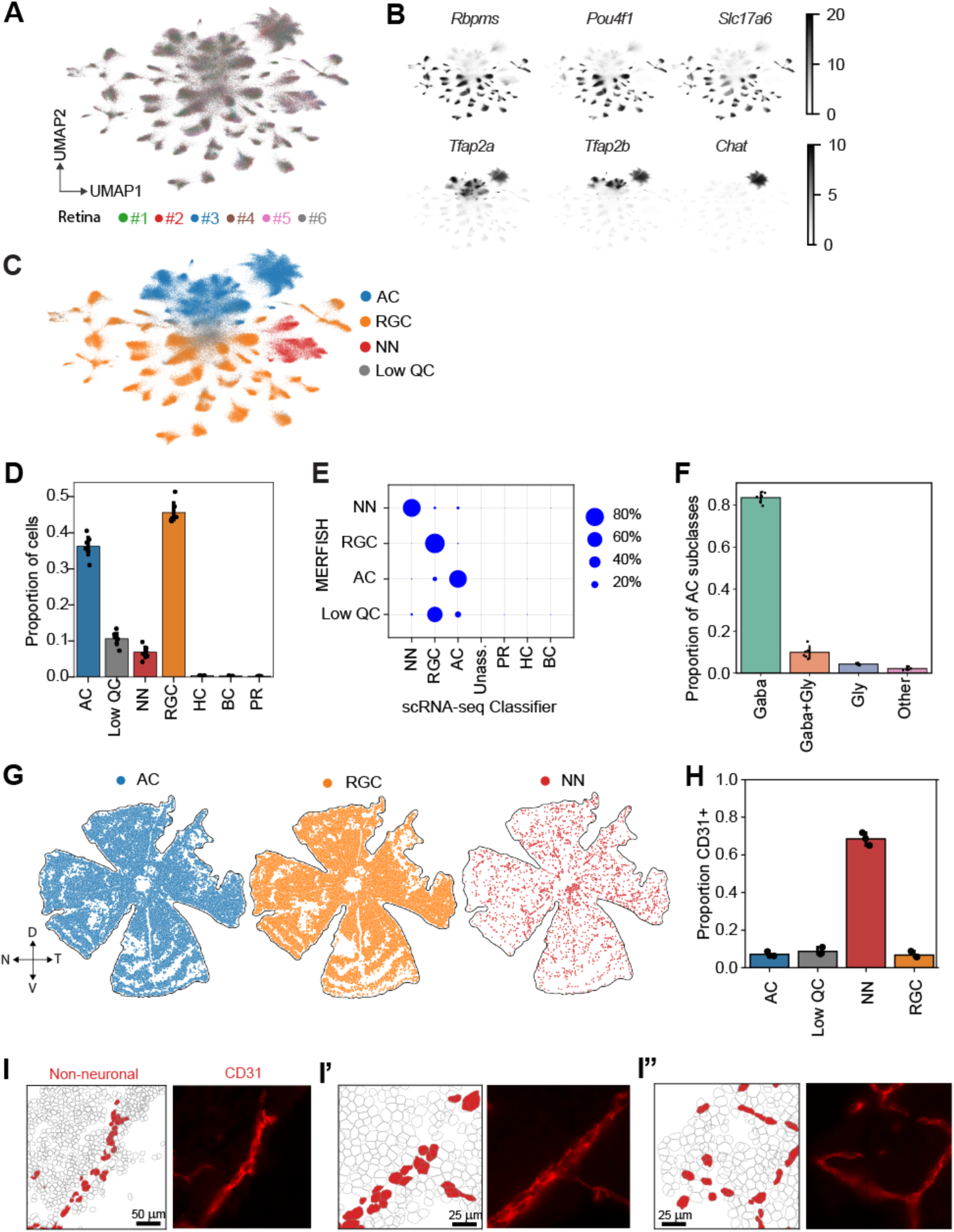
MERFISH-based categorization of retinal ganglion cells (RGCs), displaced amacrine cells (dACs), and non-neuronal cells (NNs) in 2D flatmounts. A. 2D visualization of cellular diversity in the MERFISH datasets. Cells are colored by their sample of origin (n=6 replicates). B. Same as A, with each cell shaded by marker gene expression level. Top row panels correspond to RGC markers *Rbpms, Pou4f1,* and *Slc17a6*. Bottom row panels correspond to AC markers *Tfap2a*, *Tfap2b*, and *Chat*. Note that while all ACs express *Tfap2a* and *Tfap2b*, *Chat* marks ON-starburst amacrine cells (ON-SACs). C. Same as A, with cells colored by their class of origin. RGCs and dACs are colored orange and blue, respectively, based on panel B. Non-neuronal cells (NNs, red) were identified using the XGBoost classifier (see **Figure S2C**). Low-quality cells (11%) are shown in grey. D. Quantification of RGC, AC, and NN proportions across replicates (dots). E. Confusion matrix comparing cell class assignments based on marker genes and XGBoost classification. Each row is normalized to add to a total of 100%. Notably, there are no assignments to horizontal cells (HCs), Photoreceptors (PRs), and bipolar cells (BCs), which are absent in the GCL. F. Composition of dAC subtypes grouped by neurotransmitter profiles, showing dominance of GABAergic ACs (n=6 replicates). G. Example retina displayed three times, highlighting RGCs (orange), ACs (blue), and the NNs (red) with axes indicating orientation. H. Proportions for each cell class categorized as CD31^+^ cells in MERFISH samples confirm the reliable assignment of CD31^+^ cells onto blood vessels (mean ± SD; n=6 replicates). I. Three regions (I, I’, I’’) highlighting blood vessels that can be traced using MERFISH. In each case, the left panel highlights NNs (red) among segmented cells, and the right panel shows immunostaining for CD31, an endothelial cell marker, in the same region.

dACs are axon-less neurons in the GCL composed of several distinct types (Choi et al., 2023b; Yan et al., 2020a). Consistent with their known neurotransmitter profile, ∼85% of dACs in our dataset were classified as GABAergic (Figure 2F). Notably, 32 ± 3% of dACs expressed *Chat,* the gene encoding choline acetyltransferase, identifying them as ON starburst amacrine cells (ON-SACs), the most abundant dAC type (Perez De Sevilla Muller et al., 2007) (Figure 2B). ON-SACs innervate direction-selective ganglion cells (DSGCs), including both ON-OFF DSGCs and ON DSGCs (Hamilton et al., 2021; Wei and Feller, 2011). However, because our gene panel was optimized for RGC classification, a detailed analysis of dAC diversity in our dataset was not feasible.

NNs comprised 4,206 ± 942 cells per retina, including pericytes, microglia, astrocytes, and endothelial cells. While our RGC-focused gene panel precluded finer subtyping of these NNs, their spatial distribution was distinct from that of RGCs and dACs and could be used to trace blood vessels in the GCL (Figure 2G). To confirm the identity of these vessels, we modified our MERFISH protocol to co-detect immunohistochemical signals along with RNA transcripts and cell segmentation (**Methods**). Specifically, we targeted CD31, an endothelial cell marker, to confirm the presence of blood vessels predicted computationally (Figures 2H**, 2I)**. Moreover, ∼70% of NNs in our dataset are CD31^+^, identifying them as endothelial cells. Having established the separation of major cell classes in the GCL, we now focus on dissecting the molecular diversity within RGCs using the MERFISH data.

### Consistent profiles and frequencies of RGC types between MERFISH and scRNA-seq data

*De novo* clustering based on the measured gene panel identified 35 RGC clusters (Figure 3A, **Methods**). We applied a supervised classification approach to assign adult RGC-type identities to these clusters, training the classifier on our previous scRNA-seq atlas (Tran et al., 2019), using only the genes present in the current MERFISH panel as features. The scRNA-seq atlas comprises 45 RGC clusters (C1-C45), approximately half of which have been previously shown to map 1:1 to RGC types defined by morphology and physiology (Goetz et al., 2022; Huang et al., 2022; Tran et al., 2019).

**Figure 3.**
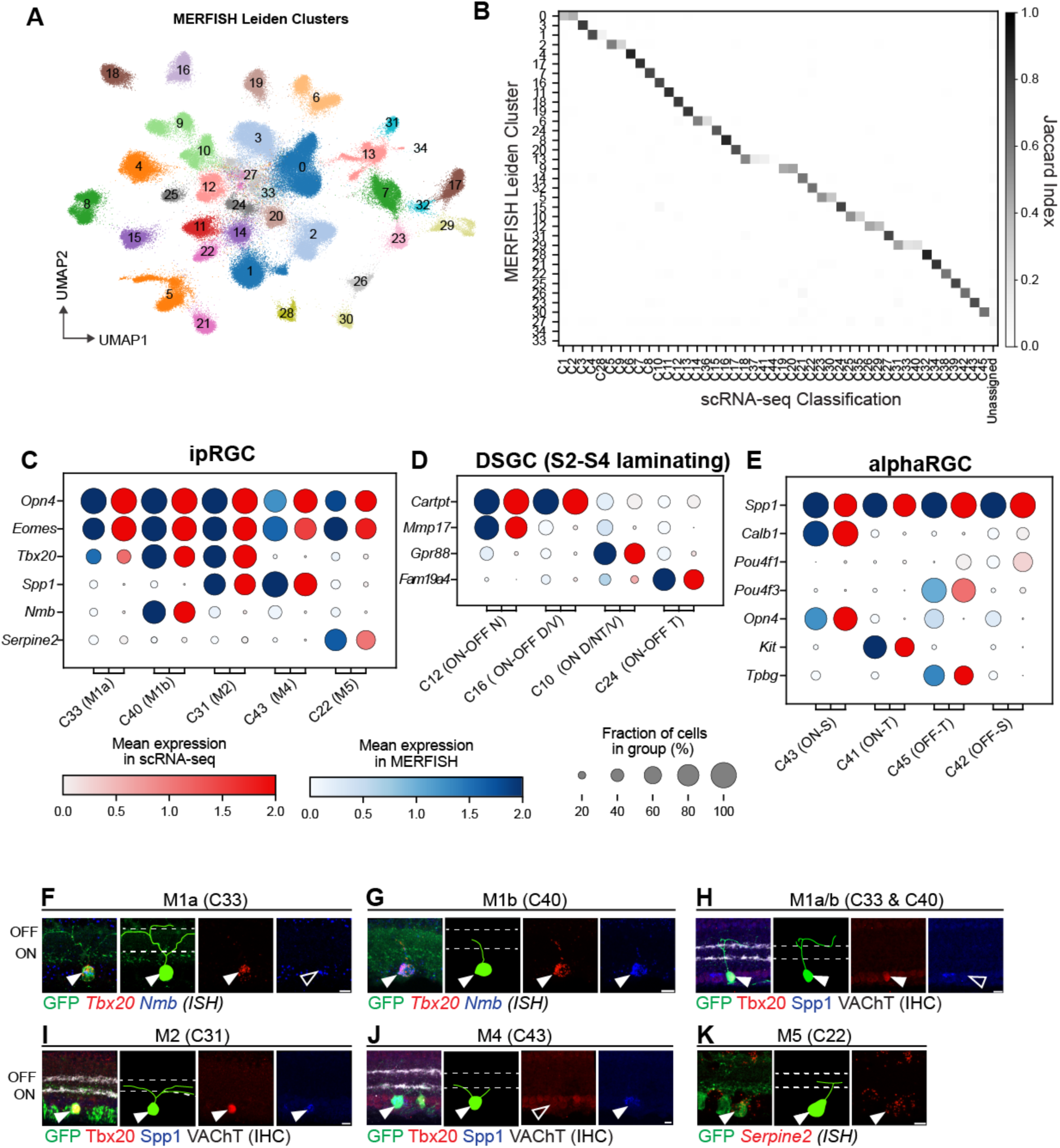
MERFISH-based RGC type classification and histological validation of ipRGC types. A. UMAP visualization of 35 MERFISH-based RGC clusters. B. Confusion matrix showing transcriptomic correspondence between MERFISH clusters 1-35 (rows) and RGC types C1-C45 (columns) defined previously using scRNA-seq (Tran et al., 2019). Darker grey shading indicates higher correspondence, quantified by the Jaccard Index. C-E. Dotplots comparing the transcriptomic fingerprints for three RGC subclasses between MERFISH (red) and scRNA-seq (blue), including clusters corresponding to *Spp1*^+^ αRGC types (panel C), *Opn4*^+^ ipRGC types (panel D); and DSGC types (panel E). The size of each dot corresponds to the fraction of cells in each group expressing the gene, and the color is the average expression level. Note that in panel E, C16 contains both dorsal (D)- and ventral (V)-preferring ON-OFF DSGCs, while C10 contains D, V, and nasal-temporal (NT)-preferring ON DSGCs. F-G. Fluorescent *in situ* hybridization (ISH) using the Opn4-Cre; LSL-YFP line for ipRGC clusters (Ecker et al., 2010), C33 and C40, corresponding to M1a and M1b. *Tbx20^+^Nmb*^-^YFP^+^ C33 (M1a) RGCs (panel F) and *Tbx20^+^Nmb*^+^ YFP^+^ C40 (M1b) RGCs (panel G) exhibit OFF dendrites characteristic of M1 ipRGCs. H. Immunohistochemistry (IHC) experiments show that Tbx20^+^ Spp1^-^YFP^+^ RGCs (M1a/b) are OFF laminating. I-J. Same as H, showing that Tbx20^+^Spp1^+^YFP^+^ C31 (M2) RGCs are ON laminating with small soma size (panel I), while Tbx20^-^Spp1^+^ YFP^+^ C43 (M4) are ON laminating with large soma size (panel J). K. ISH showing that *Serpine2*^+^ YFP^+^ C22 (M5) RGCs are ON laminating. Scale bars in panels F-K, 5 μm.

Small sets of RGC types sharing functional, molecular, and morphological features have been termed subclasses. Prominent subclasses include the DSGCs, comprising three ON types and four ON-OFF types (Hamilton et al., 2021); six types of ipRGCs, whose expression of Melanopsin (Opn4) enables them to detect visual inputs directly independent of rods and cones (Ecker et al., 2010; Quattrochi et al., 2019); four types of αRGCs, identified by their characteristically large somata (Krieger et al., 2017), with α-ON-Sustained RGCs being M4 ipRGCs (see below); and five types of *Foxp2^+^* F-RGCs (Rousso et al., 2016). **Table S2** summarizes these subclasses, stratifying the types within each subclass based on the current knowledge of their combined transcriptomic, morphological, and functional identities.

Overall, the correspondences between MERFISH clusters and transcriptomic types were highly specific (Figures 3B**, S3A)**: Of the 35 MERFISH clusters, 24 mapped 1:1 to a single RGC type, 8 clusters mapped to 2-3 types, and 3 could not be classified due to low transcriptomic quality but possess RGC markers. As three examples of multi-mapping: (1) MERFISH cluster 29 contained scRNA-seq types C31, C33, and C40, which correspond to *Opn4*^+^ ipRGC types M1 and M2 (see below), (2) MERFISH cluster 1 mapped to C4 and C28, corresponding to the F-RGC types, F-mini-OFF and F-midi-OFF respectively (Rousso et al., 2016), and (3) MERFISH cluster 8 contained the dorsal (D) and ventral (V) responsive ON-OFF DSGCs (Duan et al., 2018), which are both present in the scRNA-seq type C16, but can be distinguished by the markers *Calb1* and *Calb2* (Figures 3A**, S3A, S3B**). A detailed examination suggested that all cases of multi-mapping involved the co-clustering of closely related types, which could be further resolved using supervised analyses.

Specifically, the classifier enabled us to transfer scRNA-seq cluster labels (C1-C45) onto the MERFISH dataset. As a *post hoc* assessment of the label transfer, we examined the gene expression patterns and relative frequencies of matched RGC types between MERFISH and scRNA-seq. Figures 3C-3E and **S3C** show concordant gene expression fingerprints for four RGC subclasses: ipRGCs, ON-OFF DSGCs, αRGCs, and F-RGCs, respectively. Notably, these comparisons also revealed that a small subset (10/140) of the MERFISH probes were not effective in detecting transcripts (**Figure S3D**). Nonetheless, the label transfer procedure was robust to these probe failures due to its reliance on multi-gene patterns, as excluding these probes only slightly improved the results. We also found that relative frequencies of the 45 RGC types were highly consistent across MERFISH replicates and correlated well with scRNA-seq frequencies (Tran et al., 2019) (**Figure S3E** and **Table S2**). Finally, we also estimated soma volumes for RGC types from MERFISH images and compared them with prior estimates from electron microscopy for cross-annotated types (Bae et al., 2018) (**Figures S3F, S3G**). While the rank order of the soma volumes correlated well between the two datasets, MERFISH-based soma volume estimates for large RGC types, such as αRGCs, were systematically lower than EM estimates due to segmentation challenges (**Figure S3G)**, even though αRGCs had the highest soma volumes (**Methods)**. By contrast, soma volume estimates in MERFISH were accurate for smaller RGC types, such as F-RGCs (Rousso et al., 2016) (**Figure S3G**). Overall, these results show that the quality of RGC classification in MERFISH aligns favorably with existing transcriptomics and morphology-based classification data.

### Annotation of intrinsically photosensitive (ip) RGC types

We leveraged these results to match molecular signatures to morphology for ipRGC clusters, all defined by the expression of *Opn4*, the gene encoding Melanopsin (Figure 3C). Previous work has identified six ipRGC types in mice, named M1-M6 (Berry et al., 2023; Do, 2019; Quattrochi et al., 2019). Our prior scRNA-seq atlas suggested putative annotations of *Opn4^+^* clusters to ipRGC types: C33 and C40, which express the highest levels of *Opn4*, were labeled M1a and M1b; C31 as M2; C43 as M4, also known as the ON-sustained (S) αRGC; C22 as either M3 (Schmidt and Kofuji, 2011), M5, or M6. Two other clusters, C7 and C8, express low levels of *Opn4* but are positive for the transcription factor *Eomes (Tbr2)*, a molecular marker relatively restricted in the ipRGC subclass during development (Mao et al., 2014; Shekhar et al., 2022).

Although ipRGCs are defined transcriptomically based on *Opn4* expression, antibodies against Melanopsin have traditionally been noted to mark M1-M2 but not M4-M6 types without amplification (Maloney et al., 2024). Notably, M3 ipRGCs are low abundance (Berson et al., 2010; Dyer et al., 2024b; Schmidt and Kofuji, 2011), and likely not identified in our dataset. We incorporated anti-Opn4 immunostaining in our MERFISH experiments on the GCL and computed the intensity of antibody staining within each molecularly defined RGC cluster (**Methods; Figure S4A**). We found that Clusters C31, C33, and C40, which expressed the highest level of *Opn4* transcripts, also exhibited the highest intensity for anti-Opn4 immunohistochemistry (**Figures S4A-S4D**), consistent with these clusters comprising M1 and M2 ipRGCs. By performing fluorescent *in situ* hybridization (ISH) in a genetic line that marks all major ipRGCs (Opn4-Cre; LSL-YFP), we verified that both C33 (YFP^+^*Tbx20^+^Nmb^-^*) and C40 (YFP^+^*Tbx20^+^Nmb^+^*) possess dendrites that laminated in the S1 sublamina of the inner plexiform layer (IPL) (Figures 3F,3G), which are the hallmark of M1 ipRGCs. Moreover, YFP^+^*Spp1^-^Tbx20^+^*, encompassing both C33 and C40, were OFF-laminating, validating their posited identity as M1 ipRGCs (Figure 3H). Among other ipRGC clusters, C31 and C43 express *Spp1* (Figure 3C). We found that YFP^+^*Tbx20^+^Spp1^+^* RGCs (C31) and YFP^+^*Tbx20^-^ Spp1^+^* RGCs (C43) possessed S4/5 laminating dendrites, indicative of ON RGCs (Figures 3I**, 3J**).

The soma sizes of C43 RGCs were larger, consistent with these being M4 ipRGCs (ON-Sustained αRGCs). Therefore, YFP^+^*Tbx20^+^Spp1^+^*cells (C31), with smaller soma sizes, correspond to M2. *Spp1* was first characterized as a marker covering all αRGCs. As an addition, we verified that *Tbx20^+^Spp1^+^* RGCs did not co-label YFP^+^ RGCs in the αRGCs marking line (Kcng4-Cre: LSL-YFP) (Duan et al., 2015; Krieger et al., 2017) (**Figures S4E, S4F**). Finally, we found that YFP^+^*Serpine2^+^* RGCs possess S5 ON-laminating dendrites, identifying C22 as M5 ipRGCs (Figure 3K). Taken together, these results identify four ipRGC types among our transcriptomic clusters: M1 (C33, C40), M2 (C31), M4 (C43), and M5 (C22). C33 and C40 represent two M1 subtypes with C40 (M1b) exhibiting higher levels of melanopsin immunoreactivity than C33 (M1b). This still leaves open the molecular identity of M3 and M6. M3 is a rare ipRGC type with bistratified dendrites (Schmidt and Kofuji, 2011), whose transcriptomic identity remains unresolved. A recent study has proposed that C7, which expresses low levels of *Opn4*, is the M6 type (Dyer et al., 2024a).

### Topographic distributions of RGC types

Using the (*x, y*) coordinates in the MERFISH datasets, we examined the spatial distributions of each of the 45 RGC types on the retinal surface. For each RGC type, we assessed topographic biases of somas along the Dorsal-Ventral (D/V, represented vertically) and Temporal-Nasal axis (T/N, represented horizontally) axes (Figure 4A**, 4B**). By randomizing soma locations for each type in the dataset, we confirmed via simulations that unbiased distributions at all observed frequencies exhibit a mean score of zero along these axes (**Figure S5A, S5B)**. We then quantified D/V and T/N biases for each RGC type, finding these scores to be highly consistent across the six replicates (**Figures S5A, S5B**).

**Figure 4.**
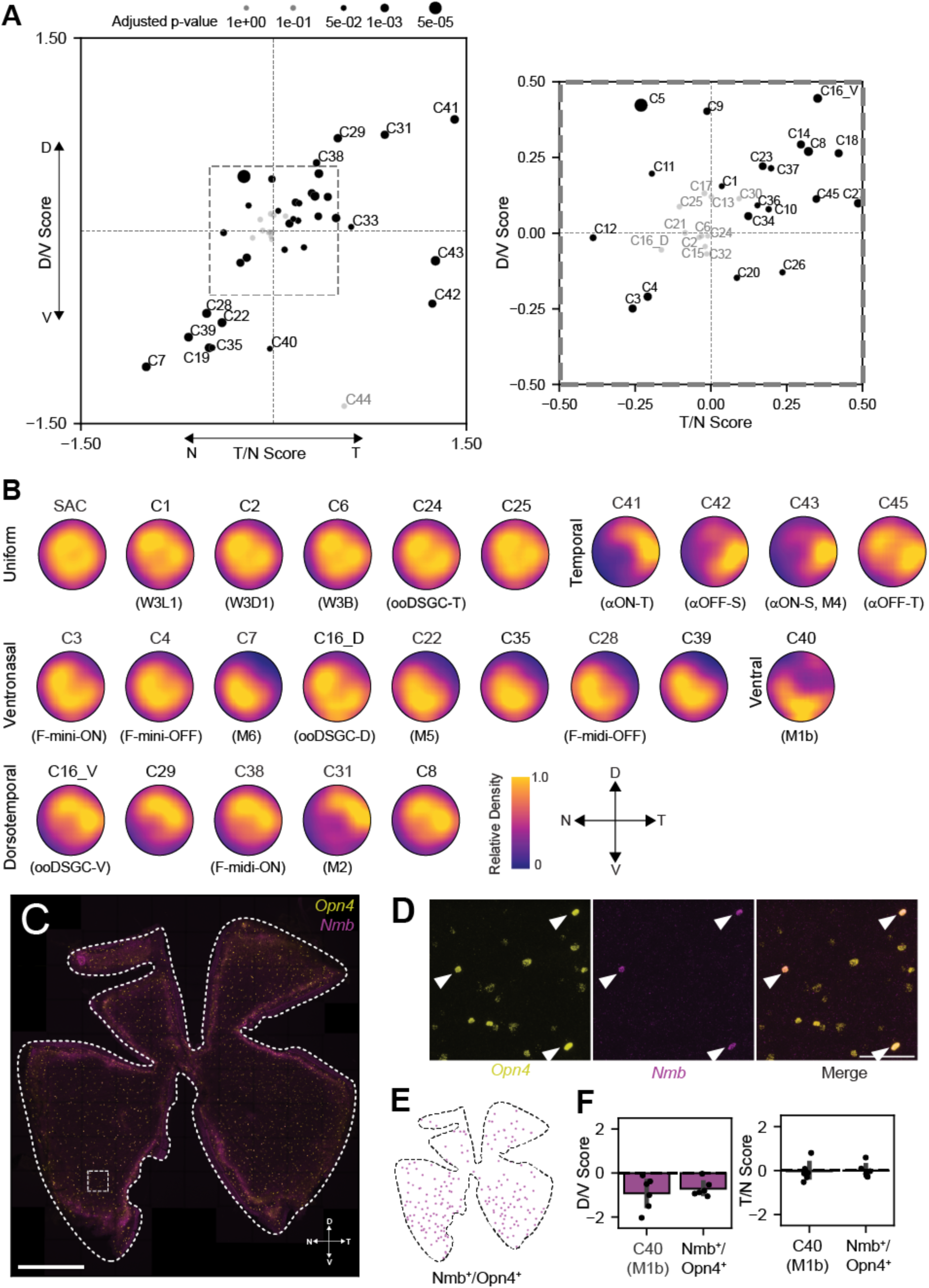
Topographic distributions of RGC types. A. Scatter plot of the Dorsal/Ventral (D/V, top to bottom, Y-axis) and Nasal/Temporal (T/N, X-axis, left to right) scores for RGC types. See **Methods** for details of the score calculations. Statistically significant deviations (Welch’s t-test, Benjamini-Hochberg correction) are highlighted in black, with larger dots for lower p-values. The inset (right) zooms in on types near the origin. Figures S5A and B show the variation in these scores across biological replicates and show that randomized somal distributions achieve a mean zero score. B. Representative RGC-type distributions grouped by uniform, temporal, ventral, or dorsal biases, with adjusted density plots accounting for sampling inhomogeneity. The distribution of Starburst Amacrine Cell (SAC) is also illustrated. **Figures S6A** and **S6B** show individual cell locations for RGC types. C-D. Wholemount ISH for *Opn4* (yellow) and *Nmb* (purple) marking C40 cells, with D/V and T/N orientations (panel C). The magnified view (panel D) shows *Nmb^+^* cells as a subset of *Opn4^+^*cells. Scale bars: 1mm in C; 100µm in D. n =6. E. Ventrally biased spatial distribution of *Nmb^+^Opn4^+^* cells in panel C. F. Comparison of D/V scores (left) and T/N scores (right) for C40 (*Nmb^+^Opn4^+^*) between MERFISH and FISH. Statistical significance was assessed using the Welch’s t-test.

Collectively, 34 out of the 45 RGC types (∼75%) showed statistically significant topographic biases along the D/V or T/N axes or both. Notably, 14 types exhibited particularly strong biases (as defined by having a D/V or T/N score exceeding ± 0.5), underscoring pervasive topographic variations among RGC types (Figures 4A**, 4B**). Intriguingly, the biases were predominantly concentrated along the dorsal-temporal (DT) and ventral-nasal (VN) axes, with a few types skewed towards the temporal quadrant (Figures 4A**, 4B**). The average topographies for types are represented in Figures 4B and **S5C,** while **Figure S6** displays some of the underlying raw data. Our findings align with previously reported topographic analyses (Heukamp et al., 2020), and report new ones. For example, three F-RGC types C3, C4, C28 corresponding to F-mini-ON, F-mini-OFF and F-midi-OFF types exhibit ventronasal biases (Figures 4C, and **S5C**), while C38 (F-midi-ON) exhibits a Dorsotemporal bias (Rousso et al., 2016). Additionally, αRGC types (C41, C42, C43, C45) displayed a Temporal bias (Figures 4C**, S5C**), though C45 (OFF-α transient) showed less bias than previously described (**Fig. S5C**) (Bleckert et al., 2014).

Among ipRGC types, several exhibited notable biases, which align with frequency estimates from previous studies (Hughes et al., 2013): M2 (C31) was dorsotemporally biased, M4 (ON-α sustained) was temporal, and M5 was ventronasally biased (Figures 4A**, 4B, S5D**). M1a exhibited a slight temporal skew, while M1b (C40) was biased ventrally. The ventral skew of M1b (C40) is consistent with a recent report showing that ventral M1 cells exhibit a higher intensity of melanopsin immunoreactivity (Hughes et al., 2013). C7, which putatively aligns with M6, is strikingly ventronasally biased (Figure 4B). To further validate our findings, we performed double fluorescent *in situ* hybridization on retinal flatmounts, focusing on C40 (*Nmb^+^Opn4^+^*) (Figures 4C**, 4D**). We confirmed that *Nmb* co-labels with *Opn4^+^* RGCs were more frequently in the ventral than in the dorsal retina, and the estimated D/V and T/N scores corroborated our MERFISH-based estimates (Figures 4E**, 4F**). Finally, the topographic distributions of M1, M2, and M4-M6 ipRGC types in our data were consistent with those identified in a recently published transgenic line (Dyer et al., 2024a) (**Figure S5D**). Overall, these findings reveal that topographic bias is a prevalent and defining feature among RGC types, with distinct spatial preferences along the DT and VN axes. Such orientation patterns may underlie functional specializations within the retina, contributing to its role in optimized visual processing.

### Regularity in the somal spacing of RGC types

A widely accepted principle of retinal organization is that each cell type forms a “mosaic” on the retinal surface (Wassle and Riemann, 1978), with somas of the same neuronal type less likely to be adjacent than expected by random chance but randomly distributed relative to somas of other neuronal types (Eglen, 2006). Mosaic arrangements are thought to ensure equitable representation of visual features across the retina, minimizing redundancy (Baden et al., 2016; Roy et al., 2021).

Although mosaicism has been extensively studied for some outer and inner retinal neuron types, such as starburst amacrine cells and cone photoreceptors (Ahnelt et al., 2000; Kay et al., 2012; Keeley et al., 2020; Raven et al., 2005), comprehensive analyses of mosaic arrangements across all RGC types using consistent criteria remain largely unexplored, with prior studies focusing only on a limited number of RGC subsets identified through genetic or histological markers (Sanes and Masland, 2015).

Using our spatial maps, we computed the nearest-neighbor regularity index (NNRI) for 33 high-frequency RGC types. We compared these to random distributions constrained by observed cell type densities, soma sizes, and positions (Kay et al., 2012; Keeley et al., 2020; Wassle et al., 1981a; Wassle et al., 1981b) (**Methods**). As a control, we confirmed that starburst amacrine cells (SACs), whose mosaic organization has been extensively documented, exhibit higher NNRIs than random (**Figure S7A**) (Kay et al., 2012; Whitney et al., 2008). Approximately 80% of tested RGC types had higher NNRIs than randomized arrangements (**Figure S7D**), consistent with mosaicism. Calculations using the Voronoi domain regularity index (Keeley et al., 2020), yielded qualitatively similar results despite some technical challenges in implementing this metric on our dataset (**Figure S7E, Methods**). Examples include types C3 and C4, which showed regular spacing consistent with prior findings (**Figure S7B, S7D**) (Rousso et al., 2016), and less well-characterized types such as C14, C25, and C26, which also exhibited non-random spacing (**Figure S7D**). A notable exception was C10, which is consistent with the fact that this cluster comprises three transcriptomically indistinguishable ON DSGC types, which showed lower regularity (Goetz et al., 2022; Hamilton et al., 2021) (**Figures S7C, S3D**). Simulations of mixed cell types confirmed that such mixtures tended to reduce NNRI scores (**Figure S7F**). In summary, our two-dimensional RGC atlas provides comprehensive analyses of global soma distributions and local regularity at RGC-type resolution, which will enable vision- and ethology-based studies.

### Uncovering Perivascular RGC types

Mammalian central neurons, including retinal neurons, are typically within 20 µm of blood vessels to ensure oxygen and nutrient supply for high metabolic demands (Wong et al., 2013). The retina contains a three-dimensional vascular lattice comprising three planar layers—superficial, middle, and deep—interconnected by penetrating vessels (Figure 5A). In our data, approximately 70% of NNs were identified as endothelial cells (Figure 2H), enabling us to trace several blood vessels within the superficial vascular layer (SL) (Figure 2I). This provided a foundation for analyzing the vascular proximity statistics of each RGC type.

**Figure 5.**
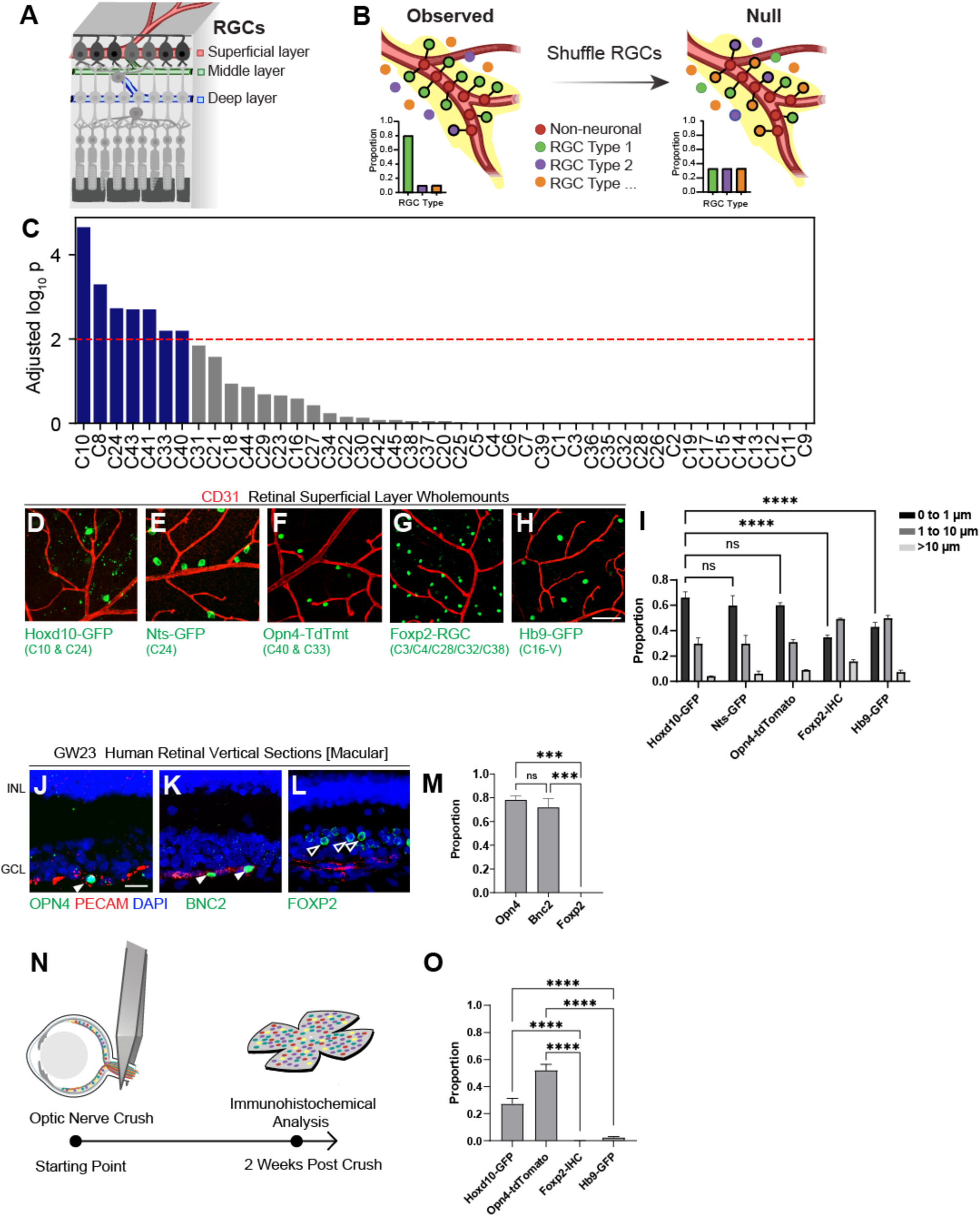
Identification and characterization of perivascular RGC types. A. Schematic showing three retinal vascular plexuses, with the superficial layer (SL) sharing the same 2D space with RGCs in the GCL. B. Permutation test schematic: random shuffling of RGC type labels generates a null distribution (right) to test the proximity of each RGC type to CD31^+^ blood vessels. C. Permutation test results highlighting that perivascular RGC types (adjusted p < 0.01) are enriched in two major RGC subclasses: ipRGCs (C40, C33, C43) and DSGCs (C24, Temporal-ON-OFF DSGCs; and C10, ON DSGCs). D-H. Validation of perivascular enrichment: Hoxd10-GFP (C10, C24 in D), Nts-GFP (C24 in E), and Opn4-TdTomato (C40, C33 in F) confirmed perivascular RGCs. In contrast, Foxp2-RGCs (G) and Hb9-RGCs (C16 in H) were not enriched in the perivascular space. Scale bars: 25µm. I: Perivascular distance measurements of RGC subclasses, grouped by proximity to vessels (0-1, 1-10, >10µm). N=4 animals each, Two-Way ANOVA tests, p<0.0001 (****). J-M. ISH in the macular region of the human prenatal retina (GW22-23) showing perivascular enrichment of *OPN4^+^* ipRGCs (J, hRGC12) and *BNC2^+^* ON DSGCs (K, hRGC11) (Yan et al., 2020b), compared to non-enriched *FOXP2^+^* F-RGCs (L, hRGC6, hRGC7). Distance for each group of human RGC subsets was quantified (M). Scale bars: 20µm. N=3 human samples each, One-Way ANOVA test, p<0.001(***). N-O. Schematic illustration of ONC experiments for mouse retina 2 weeks post crush (wpc) (N). Quantifications (O) showing that Post-ONC survival of Hoxd10-GFP (C10, C24), Opn4-TdTomato (C40, C33), at 2 weeks post crush (wpc) was higher than the average of Foxp2-RGCs (C3, C4, C28, C32, C38), or Hb9-GFP RGCs (C16), correlating with their perivascular distribution. N= 4 animals in each condition, One-Way ANOVA, p<0.0001 (****).

Using a nearest-neighbor permutation test (**Methods**), we identified 7 out of 45 RGC types that were significantly enriched for proximity to blood vessels (Figures 5B**, 5C**). 5 of these types belong to two major RGC subclasses: (1) ipRGCs – C33 (M1a), C40 (M1b), C43 (M4), and (2) DSGCs - C10 (ON DSGCs) and C24 (T ON-OFF DSGCs). Notably, we had previously identified C24 as perivascular RGCs using a candidate-based approach and revealed their role in forming the three-dimensional vascular lattice in developing retinas (Toma et al., 2024). The MERFISH-based survey not only confirms the perivascularity of C24 but also identifies the enrichment of additional RGC types within the perivascular niche.

To validate the computational results from MERFISH, we resorted to traditional transgenic labeling to characterize these perivascular RGC subsets (Duan et al., 2015; Huberman et al., 2008; Kim et al., 2010; Zhao et al., 2023). Nts-GFP, which specifically marks *Fam19a4^+^* C24 RGCs, served as a positive control based on our previous study (Toma et al., 2024). We found that 59.9±7.8% of Nts-GFP^+^ somata were located within 1 µm of CD31^+^ blood vessels (Figures 5E**, 5I**). Following this threshold, we subsequently categorized RGCs to be perivascular if they were located within 1 µm of CD31^+^ blood vessels. Next, we examined an established BAC transgenic mouse line, Hoxd10-GFP (Dhande et al., 2013), which labels subsets of ON DSGCs (C10, *Gpr88^+^*) as well as some T ON-OFF DSGCs (i.e., *Fam19a4^+^* C24 RGCs) (Figure 3D) (Dhande et al., 2013; Toma et al., 2024). In situ hybridization confirmed that *Hoxd10-GFP*^+^ RGCs consist of 54.4±2.7% *Gpr88^+^*cells (C10) and 32.5±5.6% *Fam19a4^+^* cells (C24) (**Figures S8A, B, D**). In contrast, these cells stained negative for conventional ON-OFF DSGC histology markers, including Cartpt (**Figures S8C, 8D**) (Kay et al., 2011). Notably, 66.1±4.5% of Hoxd10-GFP^+^ RGC somata were within 1 µm of CD31^+^ blood vessels (Figures 5D**, 5I**). In parallel to DSGCs, we also investigated the perivascularity of ipRGCs. Instead of the Opn4-Cre line that we utilized to mark all ipRGCs, we examined the Opn4-TdTomato BAC-transgenic line, which labels M1 (C33, C40) and M2 (C31) ipRGCs (**Figures S8E, S8F**) (Do et al., 2009), which covers the perivascular ipRGC types specifically. Our analysis showed that 60.0±2.2% of Opn4-TdTomato⁺ RGC somata were within 1 µm of CD31^+^ vessels (Figures 5F**, 5I**). As controls, we analyzed Hb9-GFP RGCs, which comprise V ON-OFF DSGCs (C16). Approximately 42.9±3.6% of Hb9-GFP⁺ RGC somata were within 1 µm of blood vessels, indicating a lower degree of perivascularity (Figures 5H**, 5I**). We also found that among Foxp2^+^ RGCs (C3, C4, C28, C32, and C38), less than 34.7±1.7% somata were within 1 µm of CD31^+^ vessels (Figures 5G**, 5I**). In summary, our results validated the MERFISH findings regarding the perivascularity of RGC types. We identified and characterized two established transgenic lines that label perivascular RGCs preferentially: Hoxd10-GFP for C10 ON DSGCs and C24 T ON-OFF DSGCs, and Opn4-TdTomato, labeling C33, C40 and C31 M1/M2 ipRGCs.

### Evolutionarily conserved perivascular RGC types in the human retina

ipRGCs and ON DSGCs are RGC types with conserved molecular profiles across mice and humans (Aranda and Schmidt, 2021; Wang et al., 2023; Yan et al., 2020b). We next examined whether these spatially restricted perivascular RGC types—such as ipRGCs and ON DSGCs—are similarly present in the human retina. OPN4 (Melanopsin) marks the transcriptomic ortholog, hRGC12 of mouse ipRGCs (M1 and M2 ipRGCs) in mice (Hahn et al., 2023; Yan et al., 2020b), while BNC2 marks hRGC11, a human ON DSGC cluster orthologous to the C10 DSGC cluster (Wang et al., 2023; Yan et al., 2020b). To explore the perivascularity of human RGC clusters, we selected OPN4 and BNC2 as candidate markers. We focused on prenatal retinas, macular regions in particular, during gestational weeks (GW) 22–23, a period of active retinal angiogenesis and neuronal growth (Lu et al., 2020). The human prenatal macular region contains multiple layers of RGCs, with blood vessels in the proximity of the bottom layer (Figures 5J-L). It gives a convenient anatomical assay to examine the perivascularity at this stage. Using such prenatal retina tissues, we detected high mRNA expression of OPN4 and BNC2 in RGCs within the GCL.78.2±3.5% of OPN4⁺ RGCs were located within 1 µm of CD31^+^ vessels (Figures 5J**, 5M**), mirroring the perivascular localization of ipRGCs in mice (Figures 5F**, 5I**). Similarly, 72.0±7.4% of BNC2⁺ RGCs were found near CD31^+^ vessels (Figures 5K**, 5M**), closely resembling the perivascular pattern observed in C10 ON DSGCs in mice (Figures 5D**, 5I**). As a control, human F-RGCs were non-perivascular and distributed away from the bottom layer of the RGCs (Figures 5L**, 5M**). Together, these results demonstrate that molecularly defined ipRGC and ON DSGC subsets are conserved perivascular RGC types in both mice and humans.

### Preferential survival of perivascular RGC subsets subject to optic nerve crush

Interestingly, the ipRGC types C33, C40, and C31, together with ON DSGCs (C10), are not only perivascularly localized but also exhibit selective resilience to axotomy among RGC types (Tran et al., 2019). This neuroprotection was previously identified through unbiased scRNA-seq analyses before and after optic nerve crush (ONC), a well-established injury model inducing systematic neuron loss (Tran et al., 2019). Independent of the transcriptomics analyses, past work of ours and others combined transgenic mouse RGC labeling and ONC also revealed that αRGCs, ipRGCs, and ON DSGCs are among the most resilient RGC subclasses subject to axotomy (Bray et al., 2019; Duan et al., 2015; Lilley et al., 2019). We performed ONC on Hoxd10-GFP and Opn4-TdTomato BAC transgenic lines (Figures 5D–5H, 5N). We noted that while αRGC resilience relies on mTOR-dependent mechanisms, as indicated by high mTOR/pS6 activity (**Figure S8I**) (Duan et al., 2015), mTOR/pS6 activities were low in Opn4-TdTomato^+^ and Hoxd10-GFP^+^ RGCs (**Figures S8G, S8H, S8J**). These data suggest that ipRGCs and ON DSGCs largely employ mTOR-independent neuroprotective strategies (Bray et al., 2019; Williams et al., 2020). We hypothesized that the perivascular localization of these RGC types (Figures 5D**, 5F, 5I**) could contribute to their resilience. To test this, we examined the survival of ipRGCs and ON DSGCs at two weeks post-ONC (2 wpc) using the Opn4-TdTomato and Hoxd10-GFP lines. Non-perivascular RGC subsets, such as Hb9-GFP RGCs and F-RGCs, served as controls. Survival rates for perivascular RGCs were significantly higher than for non-perivascular types (Figure 5O **and Figure S8K**), aligning with scRNA-seq estimates (Tran et al., 2019). Notably, axotomy did not significantly alter vascular densities at 2 wpc (**Figures S8L-N**). These findings suggest that the perivascular niche promotes RGC survival following axotomy through an mTOR/pS6 independent mechanism. Such a mechanism is an orthogonal cell-extrinsic neuroprotective strategy, distinct from those utilized in αRGCs insults (Bray et al., 2019; Duan et al., 2015; Jacobi et al., 2022; Li et al., 2022; Tran et al., 2019; Zhao et al., 2023).

## DISCUSSION

We combined MERFISH-based spatial transcriptomics with histological and computational methods to generate a comprehensive spatial atlas for mouse RGC types. By leveraging molecular signatures from scRNA-seq atlases, we registered the *en face* distributions of 45 mouse RGC types on the two-dimensional surface, providing insights into their spatial arrangement and local microenvironments. The results presented here not only validated the existing transcriptomic taxonomy of RGCs (Tran et al., 2019), but also mapped the spatial topographies of all 45 types at single-neuron resolution with minimal batch effects across adult mouse retinas.

On the technical side, our approach provides a high-throughput method to analyze RGC taxonomy in intact wild-type retinas, overcoming the limitations of previous transgenic or antibody-dependent approaches. By capturing ∼30,000 RGCs per retina, this method surpasses the throughput of scRNA-seq, which often requires pooling multiple retinas to compensate for the cell loss during enzymatic dissociation and microfluidic capture (Tran et al., 2019). Additionally, the *en face* preparation focuses specifically on the GCL, enabling precise analyses of RGC distributions, topographies, local spacing, and cell-cell interactions with the microenvironment. The flatmount approach presented here differs from a recent spatial transcriptomic analysis of vertical retinal sections (Choi et al., 2023a). As such, it can also be extended in the future to map the topography of inner retinal cell classes, such as bipolar cells and amacrine cells, in the inner nuclear layer. Additionally, our protocol also preserves RNA quality for high-resolution spatial analyses and enables the integration of antibody-based staining, broadening its applicability.

Our results reveal that over 75% of RGC types exhibit non-uniform distributions across the retinal surface, reinforcing the idea that topographic variation may be a general feature rather than an exception. (Baden et al., 2016; Heukamp et al., 2020). While our results confirm previously noted patterns for specific subsets, such as F-RGCs and αRGCs, they also suggest that this variability extends across most RGC types. This non-uniformity likely influences the structure and function of associated retinal circuits, reflecting evolutionary adaptations to the mouse’s visual environment.

Combined with our growing understanding of the evolution of retinal cell types (Hahn et al., 2023), our study presented here provides a foundation for exploring the topographic arrangements of orthologous RGC types in species with distinct visual behaviors, which will contribute to a neuroethological understanding of the vision.

By analyzing local neighborhoods, we provide evidence for regularity in cell-cell spacing among several RGC types. Moreover, we also identified multiple RGC types enriched in perivascular niches, including members of the ipRGC and DSGC subclasses. These findings, validated using two transgenic lines (Hoxd10-GFP and Opn4-TdTomato), enabled further characterization of RGC types enriched in perivascular spaces. Perivascular ipRGCs and ON DSGCs exhibited enhanced survival following axotomy compared to non-perivascular RGCs, such as F-RGCs and ventral ON-OFF DSGCs. This suggests that proximity to blood vessels may provide a neuroprotective advantage. This observation contrasts with the high-mTOR activity-dependent intrinsic neuroprotection seen in αRGCs (Duan et al., 2015; Zhao et al., 2023), pointing to distinct mechanisms mediated by the local vascular microenvironment. However, detailed mechanisms underlying neuroprotection remain to be identified. Extending this analysis to the human retina, we find that orthologous counterparts of mouse M1 ipRGCs and ON DSGCs are also perivascular in the human retina, suggesting an evolutionarily conserved relationship between these RGC types and their vascular niche. Finally, whether or not ipRGCs and ON DSGCs influence vascular patterning in the developing retina similar to C24 (T ON-OFF DSGCs) (Toma et al., 2024)) remains to be explored. The underlying neurovascular interaction mechanisms remain to be examined.

We end by acknowledging some limitations of the current work that present opportunities for future developments. The analyses focused on a curated set of 140 marker genes, limiting the ability to capture genome-wide transcriptional patterns or classify non-RGCs such as dACs and other non-neuronal types. Future advancements in spatial transcriptomics with higher gene content and capture efficiency will further expand the scope of such studies. Additionally, the use of thin tissue sections, while essential for the current approach, introduces experimental and data analysis challenges that ongoing advancements in thick-section imaging could alleviate (Wang et al., 2018). These would be especially helpful for more extensive testing of cell-cell spacing statistics, which was feasible in this dataset. Nonetheless, our work establishes a robust framework for studying RGC spatial organization, taxonomy, and interactions with the local microenvironment. This framework has implications for understanding retinal development, physiology, and neurodegeneration in mice and humans.

## ACKNOWLEDGMENTS

We thank J. He, X.K. Chen, Y. Hu, L. Li, L. Zhang, T. Puthussery, M.B. Feller, and J.R. Sanes for their advice on the study. We thank S. Varadarajan, D. Copenhagen, and K.W. Yau for sharing transgenic mouse lines. We thank Y.M. Kuo, S.L. Wang, F. Wang, E. Dang, L. Pena, P. Miller, and H. Wang for their technical support. We acknowledge NEI P30EY002162 and P30EY012196 for vision core support, RPB unrestricted fund to UCSF-Ophthalmology; from NEI (F30EY033201) to N.Y.T.; from BrightFocus Foundation Fellowship to M.Z.; from NIMH (K01MH123757) to A.T.E.; from NINDS (R35NS097305) to A.R.K; from NEI (F32EY033639) to F.C-H; from NEI (R01EY023648, R01EY034089, R01EY036071, and F01EY036430) to M.T.H.D.; from NEI (R01EY032564, R01EY027202) to B.S.; from NEI (R01EY030138) to X.D; NINDS (U01NS136405) and Glaucoma Research Foundation (Catalyst for a Cure) to K.S. and X.D; the Melza M. and Frank Theodore Barr Foundation, the McKnight Foundation, and the BrightFocus Foundation (NGR) to K.S..

## AUTHOR CONTRIBUTIONS

N.T., K.N, K.S., and X.D. conceived the study and wrote the manuscript; all authors provided input to the manuscripts; N.T., X.D. optimized histology and RNA preparation for retina MERFISH; K.N. and K.S. established computational pipeline and analyzed the MERFISH data; N.T. and M.L. performed MERFISH reactions; M.Z, M.R.L, and Y.Y performed immunohistochemistry validations; M. Z. performed animal surgeries; M.Z. and A.R.K collected human eye tissues; T.R.G and B.S. contributed to wholemount ipRGC studies; K.T., A.T.E, F.S.C-H., M.T.H.D. contributed to genetic reagents used in this study; N.R. participated in computational data analysis; K.S. and X.D. co-supervised the study and acquired research funds.

## METHODS & MATERIALS

### Contact Information for Reagent and Resource Sharing

- Computational scripts detailing MERFISH analysis reported in this paper are available at: https://github.com/shekharlab/SpatialRGC
- The processed data is in the process of being curated and will be released on Zenodo in a few weeks. The link will be posted on the above github page.
- Requests for data and code may also be directed to Karthik Shekhar; requests for genetic reagents and further inquiries may be directed to the Lead Contact, Xin Duan.

### Key resources table

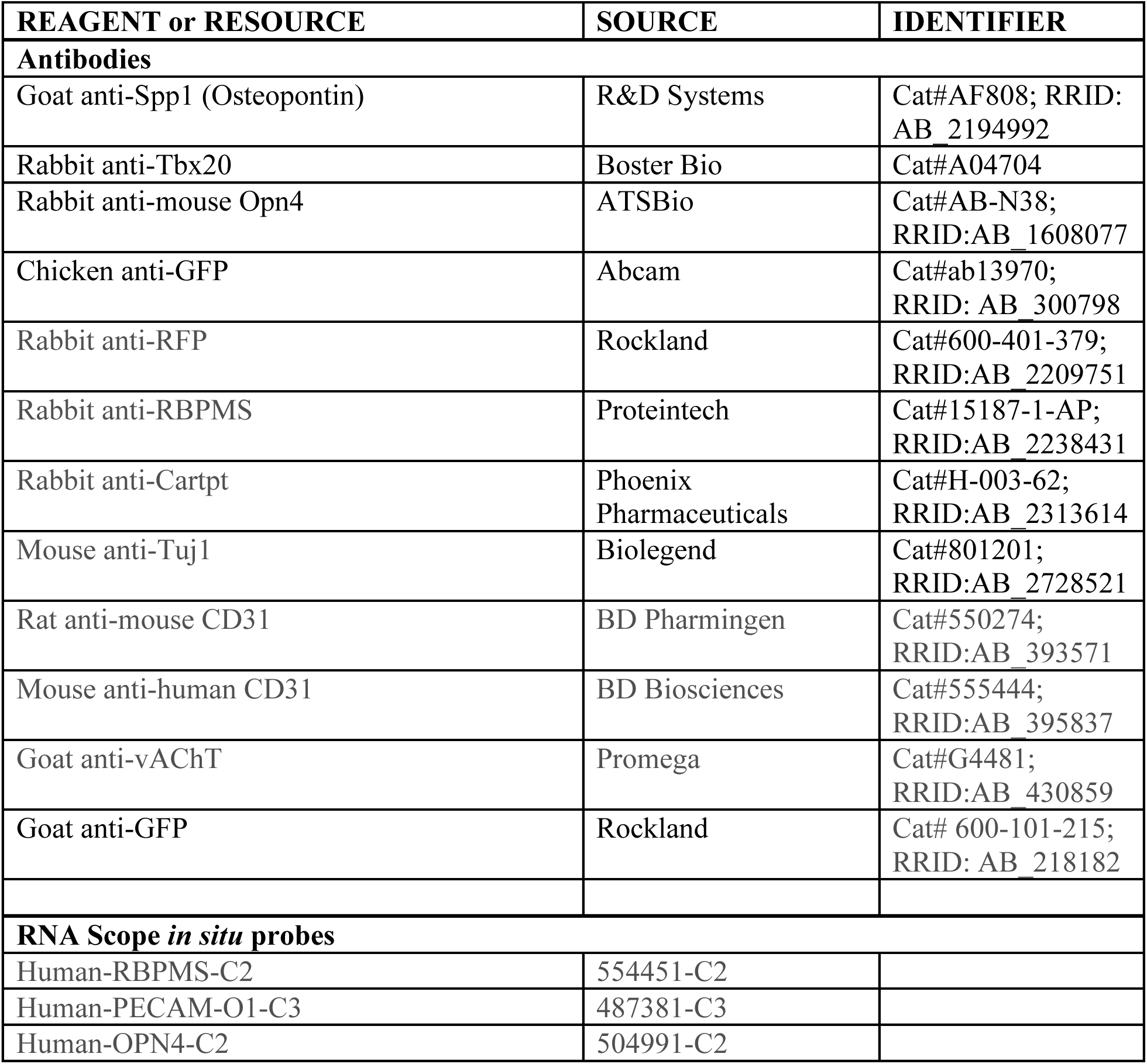
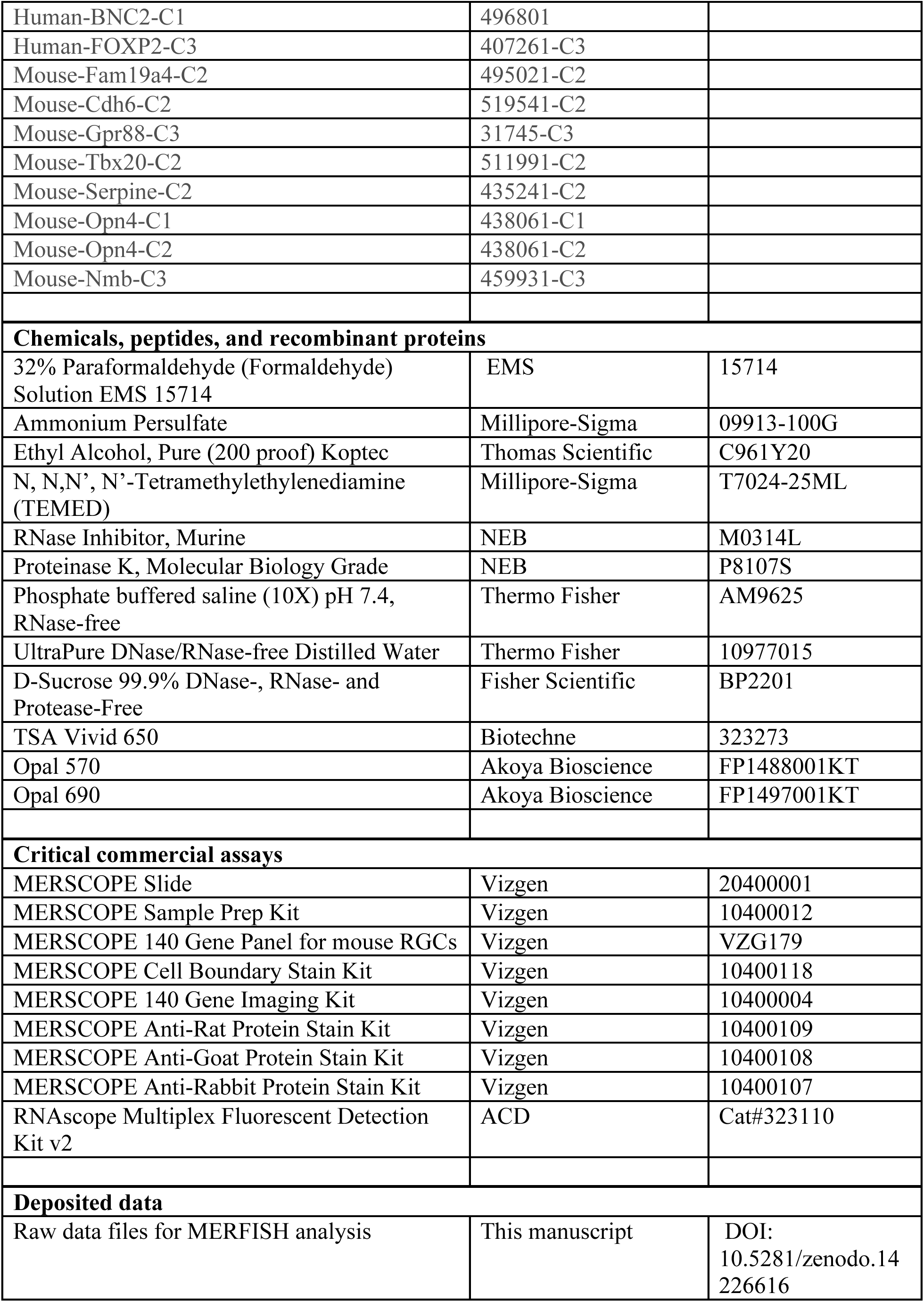
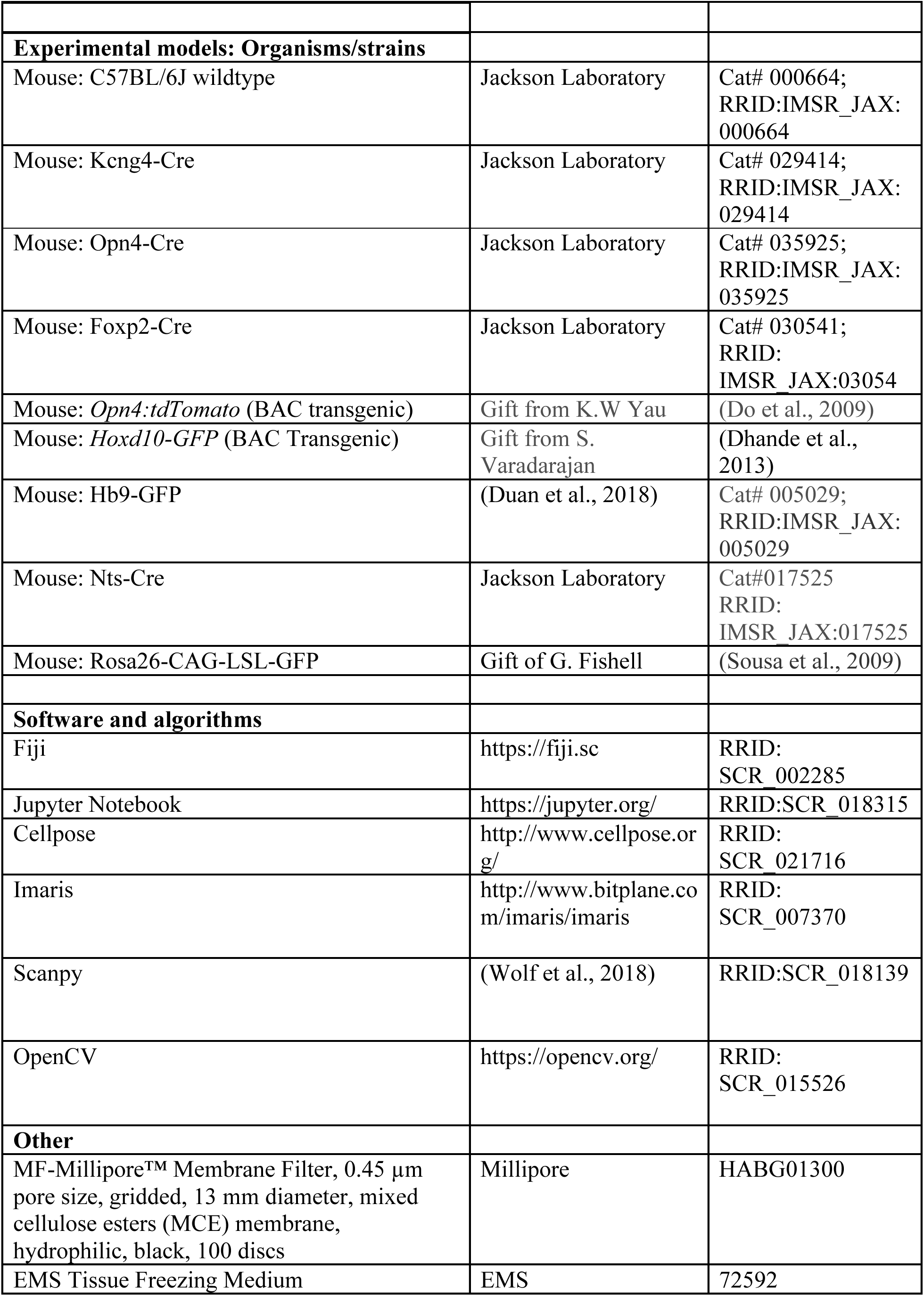
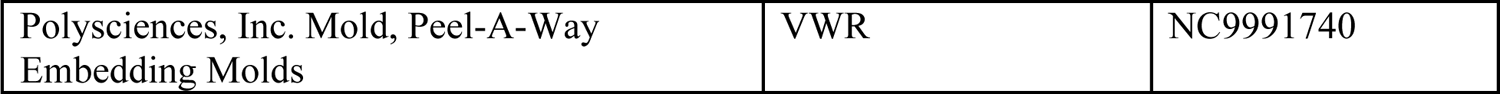

| REAGENT or RESOURCE | SOURCE | IDENTIFIER |
| --- | --- | --- |
| **Antibodies** |  |  |
| Goat anti-Spp1 (Osteopontin) | R&D Systems | Cat#AF808; RRID: AB_2194992 |
| Rabbit anti-Tbx20 | Boster Bio | Cat#A04704 |
| Rabbit anti-mouse Opn4 | ATSBio | Cat#AB-N38; RRID:AB_1608077 |
| Chicken anti-GFP | Abcam | Cat#ab13970; RRID: AB_300798 |
| Rabbit anti-RFP | Rockland | Cat#600-401-379; RRID:AB_2209751 |
| Rabbit anti-RBPMS | Proteintech | Cat#15187-1-AP; RRID:AB_2238431 |
| Rabbit anti-Cartpt | Phoenix Pharmaceuticals | Cat#H-003-62; RRID:AB_2313614 |
| Mouse anti-Tuj1 | Biolegend | Cat#801201; RRID:AB_2728521 |
| Rat anti-mouse CD31 | BD Pharmingen | Cat#550274; RRID:AB_393571 |
| Mouse anti-human CD31 | BD Biosciences | Cat#555444; RRID:AB_395837 |
| Goat anti-vAChT | Promega | Cat#G4481; RRID:AB_430859 |
| Goat anti-GFP | Rockland | Cat# 600-101-215; RRID: AB_218182 |
| **RNA Scope *in situ* probes** |  |  |
| Human-RBPMS-C2 | 554451-C2 |  |
| Human-PECAM-O1-C3 | 487381-C3 |  |
| Human-OPN4-C2 | 504991-C2 |  |
| Human-BNC2-C1 | 496801 |  |
| Human-FOXP2-C3 | 407261-C3 |  |
| Mouse-Fam19a4-C2 | 495021-C2 |  |
| Mouse-Cdh6-C2 | 519541-C2 |  |
| Mouse-Gpr88-C3 | 31745-C3 |  |
| Mouse-Tbx20-C2 | 511991-C2 |  |
| Mouse-Serpine-C2 | 435241-C2 |  |
| Mouse-Opn4-C1 | 438061-C1 |  |
| Mouse-Opn4-C2 | 438061-C2 |  |
| Mouse-Nmb-C3 | 459931-C3 |  |
| **Chemicals, peptides, and recombinant proteins** |  |  |
| 32% Paraformaldehyde (Formaldehyde) Solution EMS 15714 | EMS | 15714 |
| Ammonium Persulfate | Millipore-Sigma | 09913-100G |
| Ethyl Alcohol, Pure (200 proof) Koptec | Thomas Scientific | C961Y20 |
| N, N,N', N'-Tetramethylethylenediamine (TEMED) | Millipore-Sigma | T7024-25ML |
| RNase Inhibitor, Murine | NEB | M0314L |
| Proteinase K, Molecular Biology Grade | NEB | P8107S |
| Phosphate buffered saline (10X) pH 7.4, RNase-free | Thermo Fisher | AM9625 |
| UltraPure DNase/RNase-free Distilled Water | Thermo Fisher | 10977015 |
| D-Sucrose 99.9% DNase-, RNase- and Protease-Free | Fisher Scientific | BP2201 |
| TSA Vivid 650 | Biotechne | 323273 |
| Opal 570 | Akoya Bioscience | FP1488001KT |
| Opal 690 | Akoya Bioscience | FP1497001KT |
| **Critical commercial assays** |  |  |
| MERSCOPE Slide | Vizgen | 20400001 |
| MERSCOPE Sample Prep Kit | Vizgen | 10400012 |
| MERSCOPE 140 Gene Panel for mouse RGCs | Vizgen | VZG179 |
| MERSCOPE Cell Boundary Stain Kit | Vizgen | 10400118 |
| MERSCOPE 140 Gene Imaging Kit | Vizgen | 10400004 |
| MERSCOPE Anti-Rat Protein Stain Kit | Vizgen | 10400109 |
| MERSCOPE Anti-Goat Protein Stain Kit | Vizgen | 10400108 |
| MERSCOPE Anti-Rabbit Protein Stain Kit | Vizgen | 10400107 |
| RNAScope Multiplex Fluorescent Detection Kit v2 | ACD | Cat#323110 |
| **Deposited data** |  |  |
| Raw data files for MERFISH analysis | This manuscript | DOI:<br>10.5281/zenodo.14226616 |
| **Experimental models: Organisms/strains** |  |  |
| Mouse: C57BL/6J wildtype | Jackson Laboratory | Cat# 000664;<br>RRID:IMSR_JAX:<br>000664 |
| Mouse: Kcng4-Cre | Jackson Laboratory | Cat# 029414;<br>RRID:IMSR_JAX:<br>029414 |
| Mouse: Opn4-Cre | Jackson Laboratory | Cat# 035925;<br>RRID:IMSR_JAX:<br>035925 |
| Mouse: Foxp2-Cre | Jackson Laboratory | Cat# 030541;<br>RRID:<br>IMSR_JAX:03054 |
| Mouse: *Opn4:tdTomato* (BAC transgenic) | Gift from K.W Yau | (Do et al., 2009) |
| Mouse: *Hoxd10-GFP* (BAC Transgenic) | Gift from S.<br>Varadarajan | (Dhande et al.,<br>2013) |
| Mouse: Hb9-GFP | (Duan et al., 2018) | Cat# 005029;<br>RRID:IMSR_JAX:<br>005029 |
| Mouse: Nts-Cre | Jackson Laboratory | Cat#017525<br>RRID:<br>IMSR_JAX:017525 |
| Mouse: Rosa26-CAG-LSL-GFP | Gift of G. Fishell | (Sousa et al., 2009) |
| **Software and algorithms** |  |  |
| Fiji | https://fiji.sc | RRID:<br>SCR_002285 |
| Jupyter Notebook | https://jupyter.org/ | RRID:SCR_018315 |
| Cellpose | http://www.cellpose.org/ | RRID:<br>SCR_021716 |
| Imaris | http://www.bitplane.com/imaris/imaris | RRID:<br>SCR_007370 |
| Scanpy | (Wolf et al., 2018) | RRID:SCR_018139 |
| OpenCV | https://opencv.org/ | RRID:<br>SCR_015526 |
| **Other** |  |  |
| MF-Millipore™ Membrane Filter, 0.45 µm pore size, gridded, 13 mm diameter, mixed cellulose esters (MCE) membrane, hydrophilic, black, 100 discs | Millipore | HABG01300 |
| EMS Tissue Freezing Medium | EMS | 72592 |
| Polysciences, Inc. Mold, Peel-A-Way<br>Embedding Molds | VWR | NC9991740 |

#### Experimental Model and Method Details Mouse retinal wholemount sectioning

To track the orientation of the retina during the flatmount preparation, the nasal side of each mouse eye was marked with a marker pen. The entire globe was subsequently fixed in 4%PFA on ice for 30 min, dissected to remove the cornea and lens, and then placed back into 4%PFA on ice for another 30 min. The sclera was peeled off from the retina, and radial cuts were made to allow for flattening and orientation marking. The retina was then transferred to RNA-grade 30% sucrose/PBS and kept at 4°C until it sank. Once equilibrated, the retina was mounted onto a membrane filter (MF-Millipore). The retina and filter paper then underwent two cycles of drying and rewetting and 30% sucrose/PBS, followed by 5 min of drying. The membrane filter was trimmed to be slightly larger than the size of the retina, and the filter paper with the retina was adhered to a flat stage made from tissue freezing medium (EMS) mounted on a cryostat chuck. The retina was embedded in a thin layer of tissue-freezing medium, and the chuck was placed on a block of dry ice to solidify the tissue-freezing medium. The embedded retina was stored overnight at −80°C. Before sectioning, the retina block was equilibrated within the cryostat. Care was taken to maintain the flat orientation of the tissue while trimming and sectioning the block. Sections of 12µm thickness were collected onto MERSCOPE slides (Vizgen). Each slide was washed with DEPC-treated PBS and incubated in 70% Ethanol at 4°C overnight to permeabilize the tissue. Samples were then either processed for MERFISH imaging or stored in 70% ethanol as flatmount slides at 4°C for up to one month.

#### MERFISH workflow using MERSCOPE

The samples on MERSCOPE slides (Vizgen) were dehydrated through an ethanol series: 90% EtOH at Room Temperature (RT) for 5 minutes, 100% EtOH at RT for 1 hr, 90% EtOH at RT for 5min, and incubated with 70% EtOH at RT for 5 min. Retinal sections were then prepared for MERSCOPE imaging following Vizgen protocols for fixed frozen tissue, including the following steps as detailed in the MERSCOPE sample prep protocol:

First, Cell Boundary Staining and Protein Staining: The sample was washed with 5 mL of PBS. Blocking solution (10:1 Blocking Buffer C Premix: RNase inhibitor) was then added to the center of the tissue section and incubated at RT for 1hr. The sample was incubated in Primary Staining Solution (100:10:1 Blocking Buffer C Premix: RNase inhibitor: Primary antibody) at RT for 1 hr.

The sample was subsequently washed with 1XPBS. Secondary Staining Solution (100:10:3:1 Blocking Buffer Premix: RNase inhibitor: Cell Boundary Secondary Stain Mix: Protein Stain) was then added to the sample and incubated for 1 hr. Subsequently, the sample was washed with PBS. Fixation Buffer (4% PFA in PBS) was then added to the tissue section and incubated at RT for 15 min. The sample was then washed with PBS. In this case, we also applied the Cell boundary staining (Vizgen) for cell segmentation, as the cells within the GCL are closely packed within a monolayer, precluding DAPI and RNA-transcript-based imaging segmentation methods. (Cell Boundary #2 Channel) most clearly delineated cell boundaries in the mouse GCL and, therefore, was used for segmentation.

Second, Optimized immunohistochemistry protocols in parallel to RNA transcript detection: A set of RNAase-free antibodies was first validated, including anti-GFP (Goat), anti-Melanopsin (Rabbit), anti-Spp1 (Goat), and anti-mouse CD31 (Rat) that were used in the current MERFISH preparation.

Antibody dilutions were optimized for MERFISH experiments to ensure signal detection. Respective antibody staining kits for different species (Rat, Rabbit, Goat) were utilized accordingly and assigned to distinct imaging channels in parallel to MERFISH transcript channels.

Third, Encoding Probe Hybridization: The sample was incubated in 5 mL Formamide Wash Buffer at 37°C for 30 min. The Formamide Wash Buffer was then carefully aspirated, and the custom 140-Gene MERSCOPE Gene Panel Mix (Vizgen, VZG179) was added to the tissue section and incubated in a humidified chamber at 37°C for at least 36 hrs and a maximum of 48 hrs. Then, in the post Encoding Probe Hybridization washes, Formamide Wash Buffer was added to the sample and incubated at 47°C for 30 min. This wash step was repeated, followed by incubating the sample in Sample Prep Wash Buffer for 2 min.

Fourth, Gel Embedding: the Sample Preparation Wash buffer was aspirated from the MERSCOPE slide, and Gel Embedding Solution (2:10:1 Gel Embedding Premix:10% w/v ammonium persulfate solution:N,N,N’,N’-tetramethylethylenediamine) was added to cover the tissue sections. A coverslip pretreated with Gel Slick was placed directly over the tissue sections and incubated at RT for 90 min to solidify the gel. The coverslip was then removed. Then, the sample was treated for clearing. The sample was incubated in a pre-warmed Clearing Solution (100:1 Clearing Premix: Proteinase K) at 47°C for 24 hr.

Fifth, DAPI and PolyT staining: Following the removal of the Clearing Solution, the sample was washed with a Sample Prep Wash Buffer. DAPI and PolyT Staining Reagent were then added to the sample and incubated at RT for 15 min on a rocker. This was followed by washing with Formamide Wash Buffer for 10 min and 5 mL Sample Prep Wash Buffer.

Last, MERFISH imaging: The sample was loaded on the MERSCOPE machine for imaging, which automates the MERFISH probe exchange and readout process.

#### Gene panel simulation

To evaluate the number of genes required for RGC classification, we used three approaches as listed below. In the first approach, we identified a fixed number of differentially expressed genes for each RGC type. We began by considering all pairs of a single cluster *i* and a single gene *j* in the scRNA-seq atlas of adult RGCs (Tran et al., 2019). Next, we removed pairs wherein fewer than 40% of cells in the cluster expressed the corresponding gene. For each remaining cluster-gene (*i,j*) pair, we calculated a specificity score *S_i,j_*

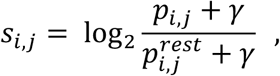

where *p_i,j_* is the proportion of cluster *i* cells expressing gene *j*, *p_i,j_^rest^* is the proportion of all remaining cells expressing gene *j,*and γ = 0.02 is a pseudocount. For each cluster *i*, we then picked the genes *j*_1_, *j*_2_ … *j*_k_ corresponding to the top *k* specificity scores *S_i,j1_*, *S*_i,j2_ … *S*_j,k_ resulting in a gene panel of size *45*k,* encompassing all 45 RGC types. Note that while a gene can be selected for more than one cluster, this did not occur in our case. We constructed gene panels corresponding to k = 1, 2, 3, and 4.

In the second approach, we randomly sampled 140 genes from the full set of ∼18k genes expressed in the data, repeating this n=10 times to estimate variance. This random gene set served as a baseline. Finally, in the third approach, we randomly sampled 140 genes from the set of 2000 highly variable genes (as selected by ‘Seurat_v3’, implemented in scanpy.pp.highly_variable_genes). As in the second approach, we repeated this n=10 times.

#### Preprocessing MERFISH data

For each 12 μm tissue section, the MERFISH protocol generated eight 1.5 μm optical sections (z_0_, z_1_, …, z_7_), spanning the ganglion cell layer (GCL) to the inner plexiform layer(IPL) along the retina’s depth. Due to the large image sizes, each optical section was scaled to be 4 times smaller along both the length and width (as implemented by OpenCV’s cv2.resize function). Each optical section was then segmented using a fine-tuned *Cellpose* model (Pachitariu and Stringer, 2022) based on the pre-trained “cyto2” algorithm. Fine-tuning was performed on sixteen manually annotated regions, each containing 50-200 soma, to improve segmentation accuracy for somas with darker staining or significantly smaller or larger soma sizes than average within the sample. The fine-tuned model used the following hyperparameters: ‘mask_threshold’=0.4, ‘flow_threshold’=0.4, and a diameter of 22, with the other parameters set to default. Additionally, rare instances of tissue regions containing only non-RGC neurons (e.g., amacrine cells, Müller glia, and bipolar cells) from the inner nuclear layer, inadvertently included during sectioning, were manually excluded from downstream analysis.

Optical sections were stitched together by assessing pixel overlap between segmented cells across consecutive optical sections. Overlap was measured using the Jaccard Index (intersection over union) between cell regions in two adjacent optical sections Z_i_ and Z_i+1_. A cell from Z_i+1_ was assigned to a cell in Z_i_ if the Jaccard index exceeded 0.15. In case of multiple cells exceeding this threshold, we picked the one with the highest overlap. Cells without sufficient overlap were treated as distinct. This stitching approach accommodated the fact that many GCL somas are less than 12.5 μm in diameter and thus only partially captured across sections. Transcripts were then registered to segmentation masks according to their pixel locations, generating a cell-by-gene matrix for each tissue section.

Soma volumes were estimated by calculating the maximal cross-sectional area of the cell across all optical sections, inferring a radius, and computing the corresponding volume. As an alternative approach to stitching, a maximum Z-projection approach was performed over optical sections, followed by segmentation. However, this approach proved unsuitable due to cell overlap distorting boundaries in the projected image. Fine-tuning the ‘cyto2’ model in *Cellpose,* particularly with 5 regions of interest (containing ∼150 manually labeled cells), significantly improved the segmentation of somas with unusual staining patterns, including darkly stained cells and smaller cells. After segmentation and stitching, tissue sections were manually aligned using DAPI-stained optical sections for rotation and translation adjustments.

For each retina, cells with fewer than 15 transcripts were excluded, and those with more than 2 transcripts of *Chat, Tfap2a,* or *Tfap2b* were labeled as expressing amacrine cell (ACs) markers. Cells meeting neither of these criteria but with more than 10 transcripts of RGC markers (Rbpms, Slc17a1, Pou4f1, Pou4f2, or Slc17a6) were considered to express RGC markers. Clustering was performed using the Leiden algorithm in Scanpy (Traag et al., 2019), and clusters were annotated as RGCs or dACs based on their dominant marker expression. About 15% of cells could not be definitively annotated and consisted of non-neuronal cells (NNs) and low-quality cells (see below).

### GCL cell class verification via XGBoost

To verify cell class assignments, an XGBoost classifier (Chen and Guestrin, 2016) was trained on published scRNA-seq datasets encompassing all retinal cell classes(Benhar et al., 2023; Tran et al., 2019; Yan et al., 2020a). Using the 130 genes that comprised the MERFISH panel (i.e., excluding the 9 failed probes in **Figure S3D** and 1 probe that was incorrectly picked during panel design), raw counts were normalized in each cell to the median value across all cells, followed by log-transformation and scaling to unit variance and zero mean. The classifier used a multiclass soft probability objective function with log-loss evaluation, 200 boosting rounds, a maximum tree depth of 4, a 40% subsampling rate, and a learning rate of 0.2. We used 30% of the dataset for validating the classifier, with rare RGC types (< 100 cells) upsampled to 100 cells and highly abundant types (>2000 cells) randomly downsampled to 2000 cells to form the training set. Cells with classification probabilities >0.5 were deterministically assigned to a type; others were marked as unassigned. The validation accuracy of this model and concordance with MERFISH clustering are displayed in **Figures S1A-C and** Figure 3B. In particular, the predictions for the model were used to annotate non-neuronal cells.

### Retinal Ganglion Cell classification

RGC types in MERFISH data were identified by classifying cells using the XGBoost model trained on the scRNA-seq atlas. The XGBoost hyper-parameters, training and validation procedure, and assignment thresholds are the same as described above. Of the 140 genes in the MERFISH marker panel, nine showed no significant expression differences compared to scRNA-seq, and one was incorrectly included.

### RGC spatial distributions in the 2D space

For each RGC type *i,* we calculated spatial bias scores along the D/V (Dorsal/Ventral) and T/N (Temporal/Nasal) axes. The D/V bias score is,

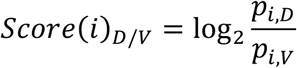

where *p*_i,+_ and *p*_j,-_ are the proportions of somas of RGC type *i* in the dorsal and ventral regions of the retina, respectively. The expression for the T/N bias score is analogous. Welch’s t-tests with Benjamini-Hochberg corrections were used to identify significant spatial biases, evaluating the null hypothesis that *Score*(*i*)_D/V_ = *Score*(*i*)_T/N_ = 0.

### Perivascularity of RGC types

To assess whether specific RGC types concentrate near retinal blood vessels, we adapted a permutation test described in a previous study (Zhang et al., 2021). Briefly, we computed a nearest-neighbor graph (n=15) connecting non-neuronal cells and RGCs, with edges removed if they were greater than 25 μm away (to account for edge effects and holes in the retina). RGC-type labels were shuffled within the same tissue section and quadrant to generate null distributions, leaving the vessel locations intact. To minimize potential global biases in the distribution, we only allowed labels to be permuted between RGCs in the same tissue section, as well as the same quadrant of the retina. We then calculated a z-score between each cell type pair (*i, j*) for a given tissue section quadrant *q*, as follows:

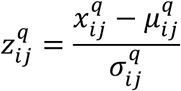

Where *X*^q^_ij_ is the observed edge count between cell types *i* and *j* in the quadrant *q*, μ^q^_ij_ is the mean number of total edges between cell types *i* and *j* in the quadrant *q* averaged across 1000 permutations, and σ^q^_ij_ is the standard deviation of the total number of edges between cell types *i* and *j* in quadrant *q* averaged across 1000 permutations. Z-scores were aggregated across quadrants using Stouffer’s method.

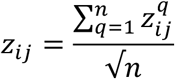

and the *z*_ij_ were used to calculate two-sided p-values *p*_ij_, under the assumption of normality

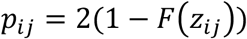

where *F* is the cumulative distribution function of the standard normal distribution. The nominal p-values *p*_ij_ were adjusted using the Benjamini-Hochberg procedure.

Density heatmaps for each type are calculated as follows. First, cells from each retina are manually aligned to the same coordinate system (x=0,y=0 is the center for each retina). Cells are assigned to discretized spatial bins of size approximately 40 μm x 40 μm each, with 101 x 101 bins total. For a given type in a given bin, a density is calculated as the number of cells of a given type in a bin divided by the total number of cells in the bin. Secondly, the relative densities for a given type are then smoothed via a Gaussian with kernel size of (41, 41) and σ_3_ = σ_4_ = 15 (implemented with cv2.GaussianBlur). ‘0’ is defined as the minimum value across bins, and ‘1’ is the maximum value across bins, with all bin values being linearly scaled accordingly.

### Mosaic analysis of each RGC type

Segmentation artifacts, cells of the same type whose cell bodies were in direct contact, were removed to avoid skewing the mosaic analysis. To identify these artifacts, we built an undirected spatial graph with vertices representing individual cells. We drew edges between vertices if they were the same cell type and within an Euclidean distance cutoff *d*. For SACs and RGC types (C41, C42, C43, C45), we chose *d*=10 μm. For all other RGC types, we chose d=4 μm. Cutoffs were chosen to be much smaller than the average diameter of the cell types to avoid removing unique cells. We then identified all connected components in the graph and kept only one node randomly from each component. This was implemented using the networkX package in Python.

For downstream mosaic analysis, nine regions of interest (ROIs) spanning 600 x 600 μm^2^ to 900 x 900 μm^2^ from different spatial locations, with 1800-3000 cells per ROI, were analyzed.

A nearest neighbor regularity index (NNRI) for each RGC type (Keeley et al., 2020; Wassle et al., 1981a; Wassle et al., 1981b). Within each ROI *r*, we calculated the distance of each RGC to the nearest neighbor of the same type *i*. The NNRI statistic for the observed distributions was calculated as

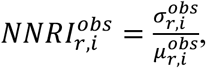

where μ_r,i_*^obs^* and σ_r,i_*^obs^* are the mean and standard deviation of nearest neighbor distances for cell type *i* in the observed ROI *r*, respectively. To generate a sample random distribution, we permuted the labels of the RGC type within each ROI n=500 times and repeated the neighbor distance calculation. The NNRI for the null distribution for a given type and ROI was then calculated as above. Note that for SACs, the null distribution was generated by shuffling labels with the other ACs.

Our second approach was to calculate the Voronoi domain regularity index (VDRI) for RGC types (Keeley et al., 2020; Wassle et al., 1981a; Wassle et al., 1981b). To generate the observed distribution in each ROI *r* for cell type *i*, we filtered out all other cell types. We drew a Voronoi diagram (utilizing *scipy.spatial.Voronoi* in Python) to obtain a distribution of Voronoi areas. Each area represents the points in space whose nearest neighbor is the contained cell. As in the nearest neighbor approach, the null distribution was created from sampling n=500 permutations with the Voronoi diagrams being redrawn. The VDRI for both the observed and null distributions was calculated analogous to the NNRI, instead using the distribution of Voronoi areas.

Here, we note some technical challenges for the Voronoi implementation. In particular, we observed that the Voronoi area distributions were artificially left-skewed due to two factors: 1) Voronoi cells near the boundaries of an ROI tend to extend excessively far (or were unbounded). 2) Voronoi cells near holes in the tissue were also larger because the cells of the same type would always be further away on the other side of the hole. To account for this, we manually drew polygons to define exclusion areas for larger holes. Any Voronoi cell that intersected the exclusion areas would have its area deducted by the area of the intersection. Similarly, Voronoi cells extending past the boundaries of the ROI would have their area truncated.

We also remark on the choice to make the null distribution a permutation of labels rather than the stochastic placement of cell bodies at random points within an ROI. This was due to 1) the non-uniform density of tissue (small holes, tissue stretching) and 2) The exclusion areas of cells, including those of amacrine cells and retinal blood vessels. Note that we report only results for RGC types with only sufficient counts of cells (corresponds to types with a proportion greater than 1.2% of total) as RGC types with too few counts produced unreliable statistics due to limited n-size.

### Genetically modified mouse lines for RGC subset labeling

All animal experiments were approved by the Institutional Animal Care and Use Committees (IACUC) at the University of California at San Francisco. Mice were maintained under regular housing conditions with standard access to drink and food in a pathogen-free facility. Male and female mice were used in equal numbers; no sexual dimorphisms were observed in retinal neuron development or vascular development, and all ages and numbers were documented. Genotypes were determined by PCR from toe or tail biopsy. Littermates were used for genetic comparison, as indicated in experimental details. Dates of birth were tracked following the vivarium birth notice and cross-referenced by JAX guidelines. Specifically, the following mouse lines were used in the following categories:

1. RGC marker lines include the following: Hb9-GFP transgenic mice express EGFP in ventral-preferring ON-OFF DSGCs (Duan et al., 2018). This transgenic line has been well characterized as to mark ON-OFF DSGCs that are Cartpt-positive.
2. Kcng4-Cre; LSL-YFP mark all αRGCs, (Duan et al., 2015; Goetz et al., 2022), including C43 (M4 ipRGC, ON-Sustained), C45 (OFF-transient), and C42 (OFF-sustained) characterized here.
3. Opn4-Cre; LSL-YFP mark M1-M6 ipRGCs (Ecker et al., 2010; Maloney et al., 2024), including M1 (C40, C33), M2 (C31), M4 (C43) and M5 (C21) ipRGCs characterized in this study. No significant M3 or M6 ipRGCs from this line were detected in the current MERFISH-based platform, which are expected given their sparseness and morphological heterogeneity (Berson et al., 2010; Schmidt and Kofuji, 2011).
4. Opn4-TdTomato (Do et al., 2009) were observed to primarily mark M1 and M2 ipRGCs (C40, C33, and C31). The Opn4-TdTomato BAC transgenic line was a gift from K.W Yau (Hopkins).
5. Hoxd10-GFP label ON DSGC subsets (C10) and a fraction of T ON-OFF DSGCs, which are Cartpt-negative (Dhande et al., 2013). We further characterized this transgenic line in **Figure S8A-D** using molecular markers based on scRNA-seq. The Hoxd10-GFP BAC transgenic line was a gift from S. Varadarajan (UT-SWMED).
6. Nts-Cre; LSL-GFP mark RGCs expressing Neurotensin (Nts), which were previously characterized to be perivascularly-enriched (Toma et al., 2024).
7. Foxp2-Cre marks RGCs in a subset of Foxp2-expressing retinal ganglion cells (F-RGCs) (Rousso et al., 2016).
8. Wildtype C57/Bl6J mice were used to carry out the set of MERFISH experiments to establish the atlas.

Animal numbers, genotypes, and ages in the current manuscript were reported in **Supplementary Table 3.**

### Immunohistochemistry for retina

Mouse eyes were collected and fixed in 4% PFA/PBS on ice for 30 minutes, followed by retina dissection, post-fixation for 30 min, and rinsing with PBS. As previously described, retinas were analyzed as cryosections and flatmounts (Duan et al., 2014). Wholemount retina samples were incubated with blocking buffer (5% normal donkey serum, 0.5% Triton-X-100 in PBS) overnight, then incubated for 2-4 days at 4°C with primary antibodies. For sectioning, fixed retinas were incubated with 30% sucrose in PBS for 2 hrs, then frozen in OCT as blocks and sectioned at 20µm. Vertical sections were incubated with 0.3% Triton X-100 and 3% donkey serum in PBS for 1 hr and then with primary antibodies overnight at 4C. Secondary antibodies were applied for 2 hours at room temperature. Retinas or sections were mounted onto glass slides using SlowFade Gold antifade reagent (Invitrogen). The following immunohistochemistry antibodies used were as follows: rabbit and chicken anti-GFP (1:1000, Millipore; 1:500, Abcam); goat anti-VAChT (1:500, Promega); rabbit anti-Cartpt (1:2500, Phoenix Pharmaceuticals); rabbit anti-Opn4 (1:1000, ATSBio); goat anti-Osteopontin/Spp1 (1:500, R&D Systems); mouse anti-Tuj1 (1:1000, Biolegend); rabbit anti-RBPMS (1:1000, Proteintech); rat anti-mouse CD31 (1:100, BD Biosciences); mouse anti-human CD31 (1:100, BD Biosciences). The primary antibodies were detected with Alexa Fluor Dye-conjugated secondary antibodies (Invitrogen). Nuclei were stained with DAPI.

### RNAscope *in situ* hybridization

*In situ* hybridization on retina sections was performed primarily using RNAscope Multiplex Fluorescent Detection Kit V2 (ACD Bio), as previously described (Matcham et al., 2024). The retina sections were post-fixed in 4% PFA and washed in PBS. After target retrieval, the sections were treated with Protease III, incubated with target probes, treated with amplification reagents, and developed with TSA Fluorescein, Cy3, and Cy5 fluorophores (Akoya Bioscience). For GFP-guided maker analysis, GFP detection by immunohistochemistry was performed after RNAscope detection. An optimized *in situ*-based GFP antibody (chicken anti-GFP, Aves Biosciences) was used for RNAscope-IHC double detection. Slides were mounted using SlowFade Gold antifade reagent (Invitrogen). RNAscope Probes included the following: RNAscope Probes: Mm-Fam19a4-C2 (495021-C2); Mm-Gpr88-C3 (317451-C3); Mm-Cdh6-C2 (519541-C2). RNAscope Probe-Hs-PECAM1-O1-C3 (487381-C3); RNAscope Probe-Hs-RBPMS-C2 (554451-C2); Human-OPN4-C2(504991-C2); Human-BNC2-C1 (496801); Human-FOXP2-C3(407261-C3).

*In situ* hybridization on retina wholemount was performed using RNAscope Multiplex Fluorescent Detection Kit v2, ACDBio, and probes described above. We slightly modified the protocols to adapt the reagents to wholemount (T.G, B.S, manuscript in preparation). Mouse Opn4 mRNA was labeled using Opal 570 (Akoya Bioscience), and Nmb mRNA was labeled using TSA Vivid 650 or Opal 690 fluorophores. All retinas were imaged on a Leica SP8 microscope at 20X magnification. Nmb-positive RGCs were counted in ImageJ/FIJI.

### Human retinal tissues

For prenatal human retina tissue collection, protocols (10-05113) were approved by the Human Gamete, Embryo, and Stem Cell Research Committee at UCSF. De-identified second-trimester human tissue samples were collected with previous patient consent in strict observance of the legal and institutional ethical regulations. Human eye tissues were processed as previously described (Zhao et al., 2023). They were harvested from eye enucleations, followed by 4% PFA fixation at 4°C overnight. Retinal tissues were removed and embedded for long-term storage and subsequent processing. GW22-23 samples were primarily used in the current study. Prenatal macular tissues were dissected into smaller ∼25 mm^2^ pieces, similar to the mouse histology. This region contains active angiongensis during development and also contains multiple layers of RGCs. The vasculature in the macular region extended into the bottom layer of the RGCs. RNA *in situ* hybridization and immunohistochemistry were carried out using the protocols established above.

### Optic nerve crush (ONC) model

ONC procedures were carried out as previously described (Duan et al., 2015). In brief, a mouse was anesthetized first, followed by a peritomy on the eye created by incising the conjunctiva at around the 4 o’clock position using a pair of spring scissors. A gentle dissection was used to bring the optic nerve into view as it exited the globe. A crush injury to the optic nerve was then made using a pair of forceps at about 2 mm from the globe for 5 sec, followed by post-op care. Contralateral eyes were used for sham control in the same animal.

### Data acquisition and quantifications for retinal immunohistochemistry data

Confocal imaging was performed using both sexes of mice. Age-matched mice were chosen for comparisons across different genetic labeling. The images were acquired using Zeiss LSM900 (Carl Zeiss Microscopy). Given the avascular regions and uneven flat mounting, imaging and quantification areas were chosen to avoid the retina’s periphery and the optic nerve head.

1. To quantify the distance from immunolabelled RGCs to the nearest vessels, images of retinal flatmounts labeling RGCs (Hoxd10-GFP RGCs, Hb9-GFP RGCs, F-RGCs, and Opn4-TdTomato) were taken using a Zeiss LSM900 microscope with a 20x objective lens with Z-stacks from the superficial layer vessels to the middle layer vessels. IMARIS was used to render the surfaces for blood vessels marked by CD31, fluorescently labeled cells, and RBPMS^+^ cells after subtracting for background noise. The “Shortest Distance to Surface” built-in function in IMARIS was used to calculate the shortest distance between RGC and blood vessel surfaces. Distances were placed into bins of 0-1 µm, 1-10µm, and greater than 10µm. RGCs within the 0-1 µm bin were defined as close contact. Comparisons between RGC types were analyzed using a two-way ANOVA with multiple comparisons test.
2. To count the contacts between human OPN4, BNC2, or FOXP2 ^+^RGCs and CD31/PECAM1^+^ blood vessels, we scored by the physical contact based on a 320 µm^2^ fluorescent ISH image set taken by Zeiss LSM900 for consistency.
3. To quantify the RGC survival under control conditions or subject to ONC treatment, we followed an established method for retina wholemount and subsequent quantification methods (Zhao et al., 2023). After ONC treatment, images were acquired using 20X magnification on a Zeiss LSM900. The survival rate was calculated by the percentage of RBPMS^+^ cells in ONC eyes against eyes that were not crushed. Retina images were acquired from at least 8 individual sections using 20X magnification. IMARIS was used to generate cell volumes after background subtraction, and the number of RBPMS^+^ cells was counted. Comparisons between the control and the 2 weeks post crush (2wpc) ONC time point were analyzed using a one-way Brown-Forysthe and Welch ANOVA test with Dunnett’s T3 multiple comparisons test.

### Statistical methods

Statistical methods and numbers of animals tested with each configuration are documented for each experiment under the figure legends. When comparing two groups, statistics were derived using an unpaired two-sided Student’s t-test or Fisher’s exact test, as indicated in the figure legends. One-way analysis of variance (ANOVA) was performed in Graphpad Prism 10 to assess the statistical significance of the differences between multiple measurements. Statistical significance was defined as p<0.05(*). p<0.01(**), p<0.001(***) and p<0.0001 (****). Statistical test values were reported in **Supplementary Table 4** for all experiments.

## SUPPLEMENTARY FIGURES

**Figure S1.**
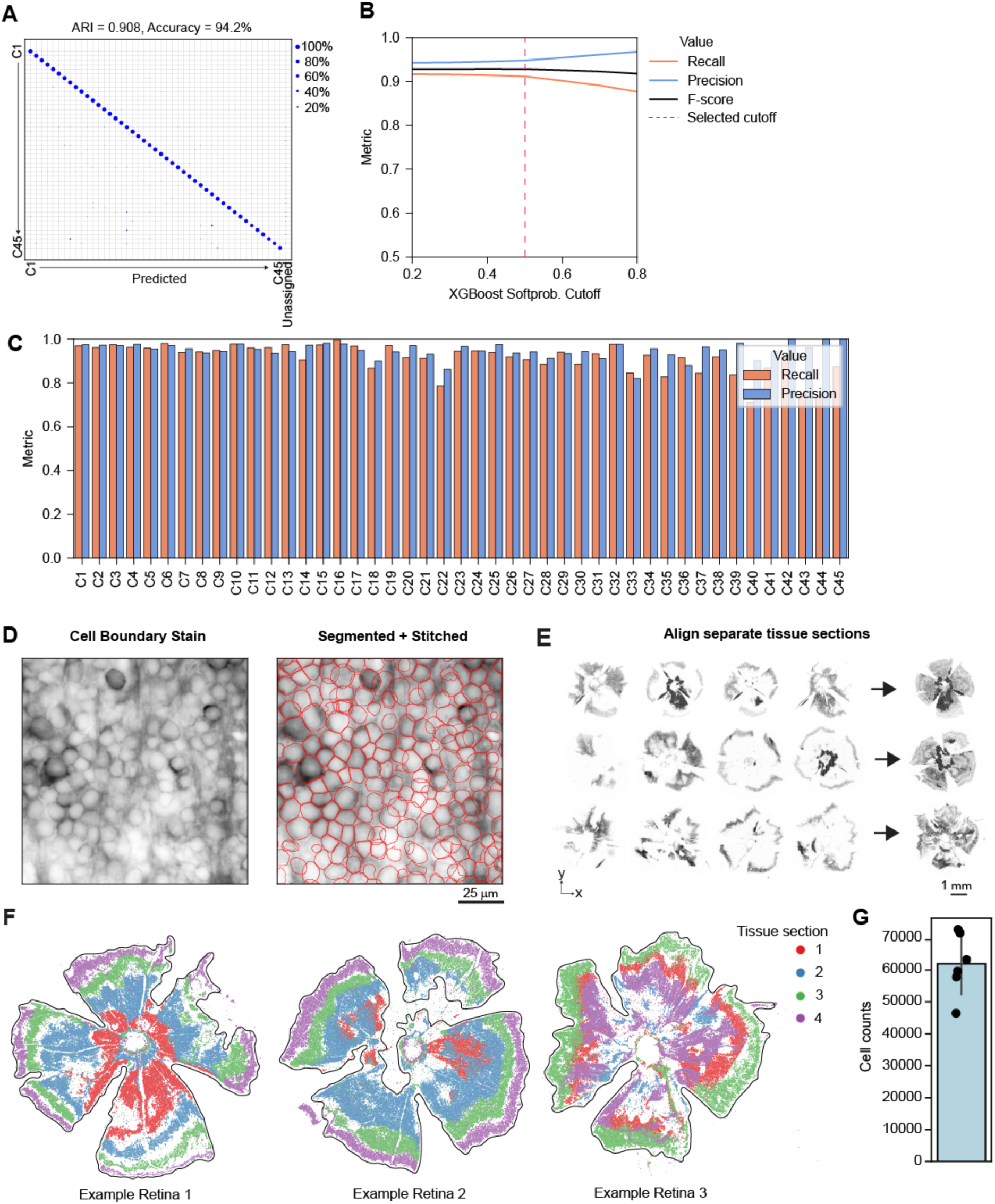
(Related to Figure 1) MERFISH gene panel selection and validation. A. Confusion matrix showing that an XGboost classifier (Chen and Guestrin, 2016) trained on the scRNA-seq atlas using the MERFISH panel as features accurately classify “held-out” RGCs (70:30 training/test split. Metrics: Adjusted Rand Index (ARI) = 0.908; Accuracy = 94.2%. B. Averaged F1-score and precision-recall metrics as a function of cutoff probability for type assignment (x-axis). Cells with a majority vote that was lower than the cutoff were labeled “Unassigned.” The vertical dashed red line indicates the value of the cutoff probability used in the study. C. Precision and recall metrics for individual RGC types for the selected cutoff probability. D. 2D image section as in Figure 1H, with the left depicting MERSCOPE cell boundary staining (grey) and the right depicting outlines of cells (red) segmented by Cellpose after stitching across multiple optical sections. E. Alignment of 4-5 tissue sections per retinal sample to reconstruct the full GCL, illustrated with three example retinae. F. Retinal cells color-coded by their originating serial sections from three retinae shown in panel (E). G. Total retinal cell counts across replicates (n=6).

**Figure S2.**
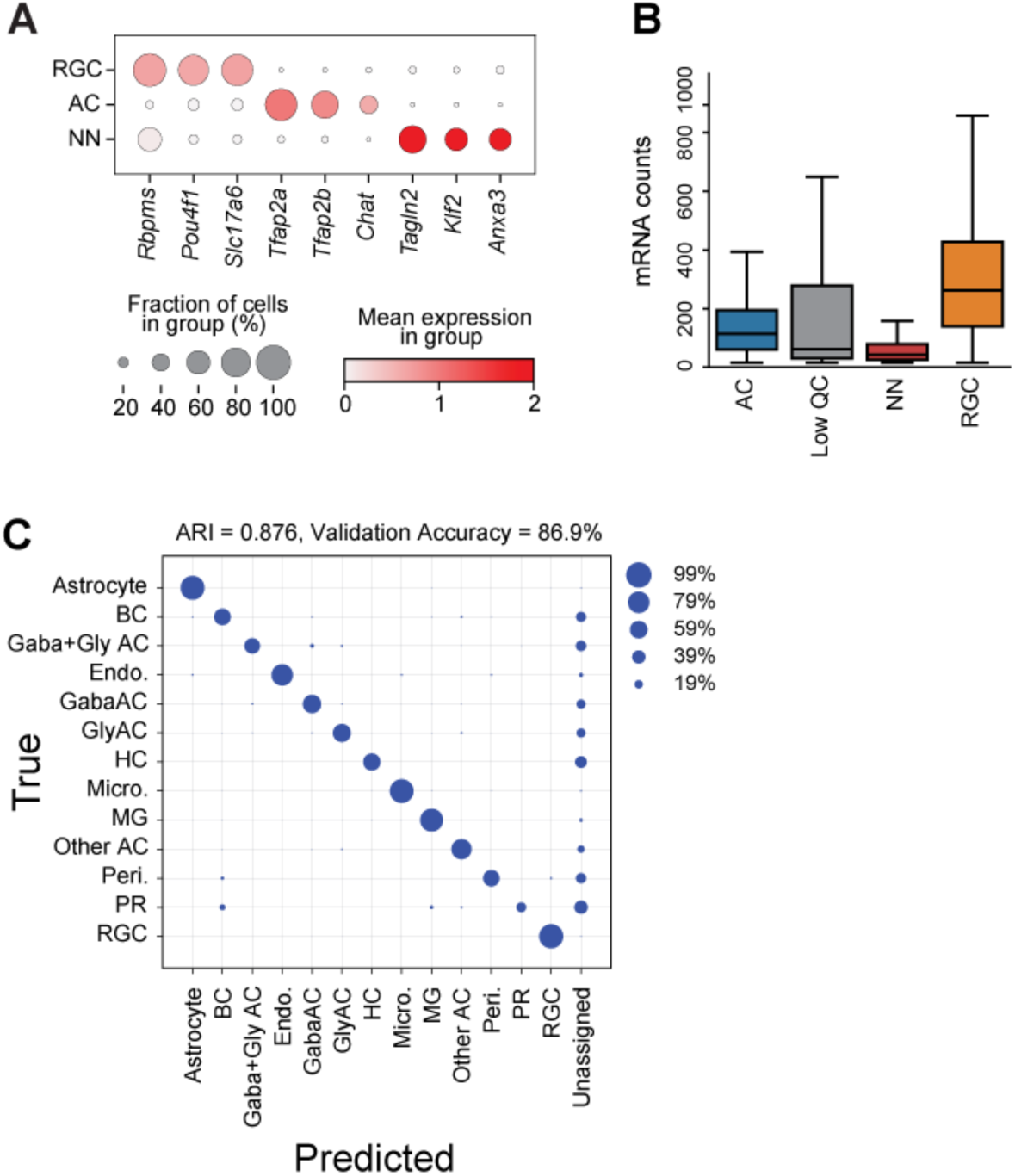
(Related to Figure 2) Additional data supporting MERFISH-based cell classification. A. Dotplot showing the expression levels of marker genes (columns) across RGC, AC, and NN classes (rows). The size of each dot corresponds to the fraction of cells in each group expressing the gene, and the color is the average expression level. B. mRNA counts per cell across four cell classes (AC, Low QC, NN, and RGC). Boxplots show the median and the interquartile range (n=6). Mean ± SD for the four groups are [AC: 141 ± 100, Low QC: 198 ± 282, RGC: 311± 229, NN: 65 ± 66]. C. Validation of the XGBoost for retinal cell classes. The classifier was trained on reference scRNA-seq data taken from three datasets (Benhar et al., 2023; Tran et al., 2019; Yan et al., 2020a), with a (70:30) for (training: validation) split (**Methods**).

**Figure S3.**
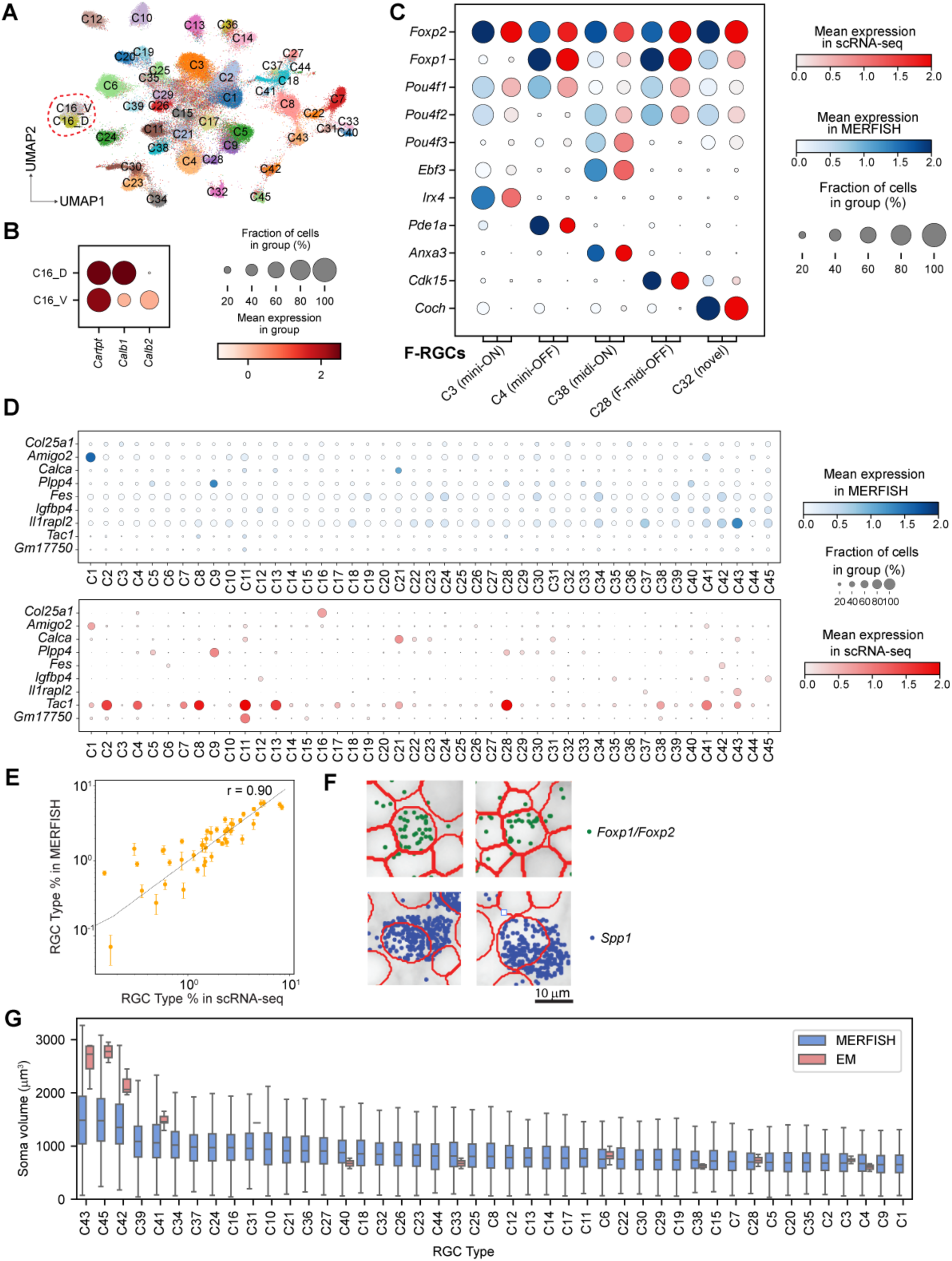
(Related to Figure 3) Additional analyses of RGC subclasses αRGCs, ipRGCs, DSGCs, and F-RGCs. A. Same as Figure 3A, with cells colored by their RGC type identity (C1-C45) based on the XGBoost classifier. C16 is split into two subtypes based on molecular markers and is circled in red (see panel B). B. Dot plot showing molecular markers *Calb1* and *Calb2* distinguishing C16-D and C16-V (ON-OFF DSGC types). Representation as in Figure S2A. C. Comparison of transcriptomic fingerprints between MERFISH and scRNA-seq for F-RGCs, defined by the expression of the transcription factor *Foxp2* (C3 corresponds to F-mini-ON, C4 to F-mini-OFF, C28 to F-midi-OFF, C38 to F-midi-ON). D. Dot-plots showing the expression patterns of 9 failed probes in MERFISH (top) compared to their expected patterns in scRNA-seq (bottom). Note that 1 additional probe not relevant to the retina was incorrectly chosen for the panel and also removed. E. Correlation (R=0.90) of RGC type proportions between MERFISH and scRNA-seq. F. Example segmentation of F-RGCs (*Foxp1^+^/Foxp2^+^*) and α-RGCs (*Spp1^+^*). Note: the under-sizing of αRGCs relative to the spatial distribution of *Spp1*. G. Soma volume distribution of selected RGC types in MERFISH (blue) compared to published EM data (pink). Note: not all RGC types were annotated at single RGC type resolution in the EM data. Refer to Table S2 for the details of all RGC types across the database.

**Figure S4.**
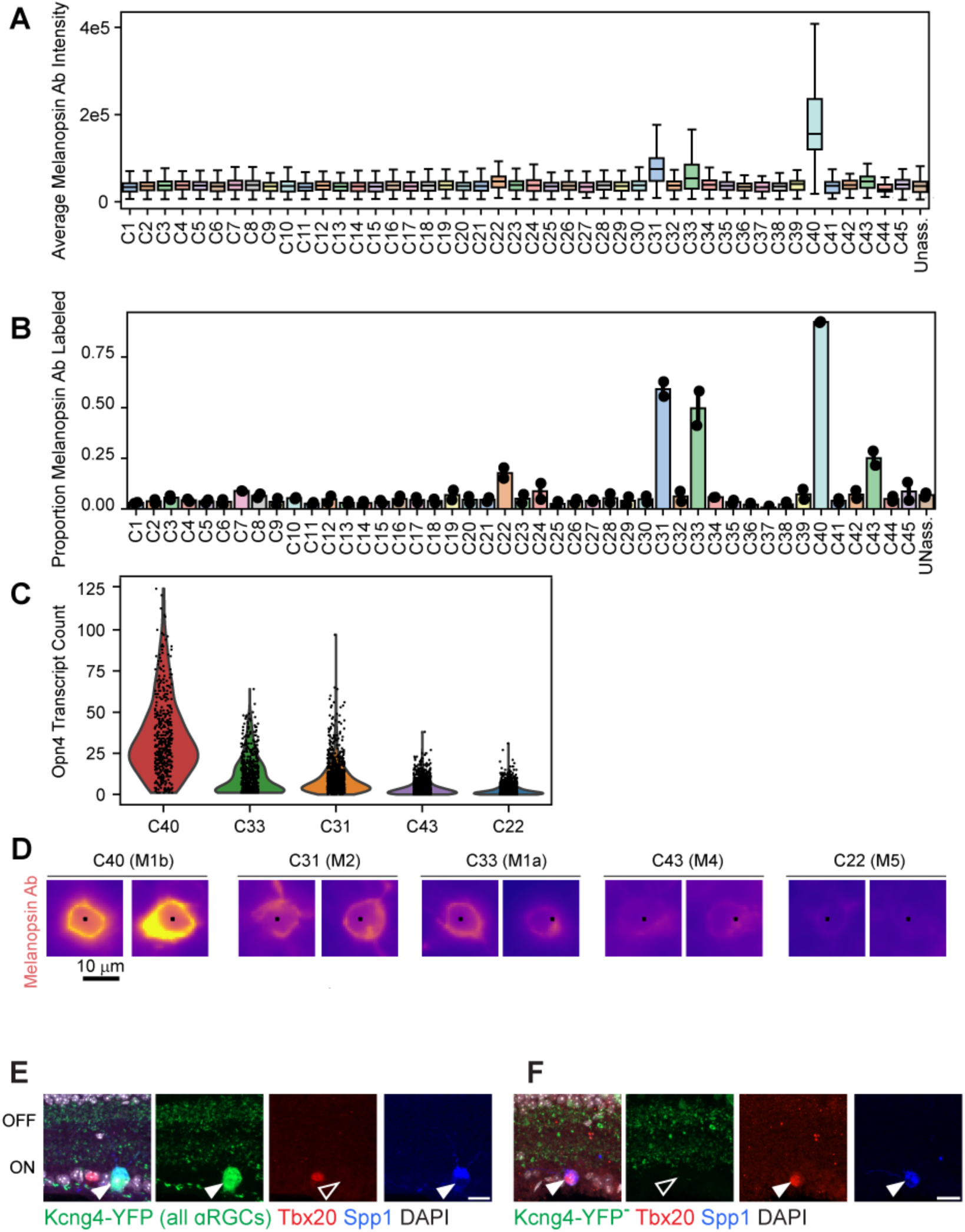
(Related to Figure 3) Additional analyses for ipRGCs in the MERFISH dataset. A. Average anti-Opn4 staining intensity across 45 RGC types. C31, C33, and C40 show high staining intensity, with C40 being the highest, identifying these clusters as M1/M2 ipRGCs (also see Figures 3F-I). B. Fractions of Opn4^+^ neurons in each RGC cluster based on thresholded staining (n=2). C. Opn4 RNA transcript counts per RGC in the five putative ipRGC clusters (n=6). D. Example images showing anti-Opn4 protein expression level across the five ipRGC clusters, with C40 expressing the highest levels (as shown in Figures S3B, 3C above). E-F. Immunohistochemistry shows that Kcng4-YFP^+^ αRGCs are Tbx20^-^Spp1^+^ (E), Tbx20^+^Spp1^+^ C31 (M2) ipRGCs are YFP^-^. Thus M2 ipRGCs are Spp1^+^, but not maked by the Kcng4-YFP line (F). Scale bars: 20µm.

**Figure S5.**
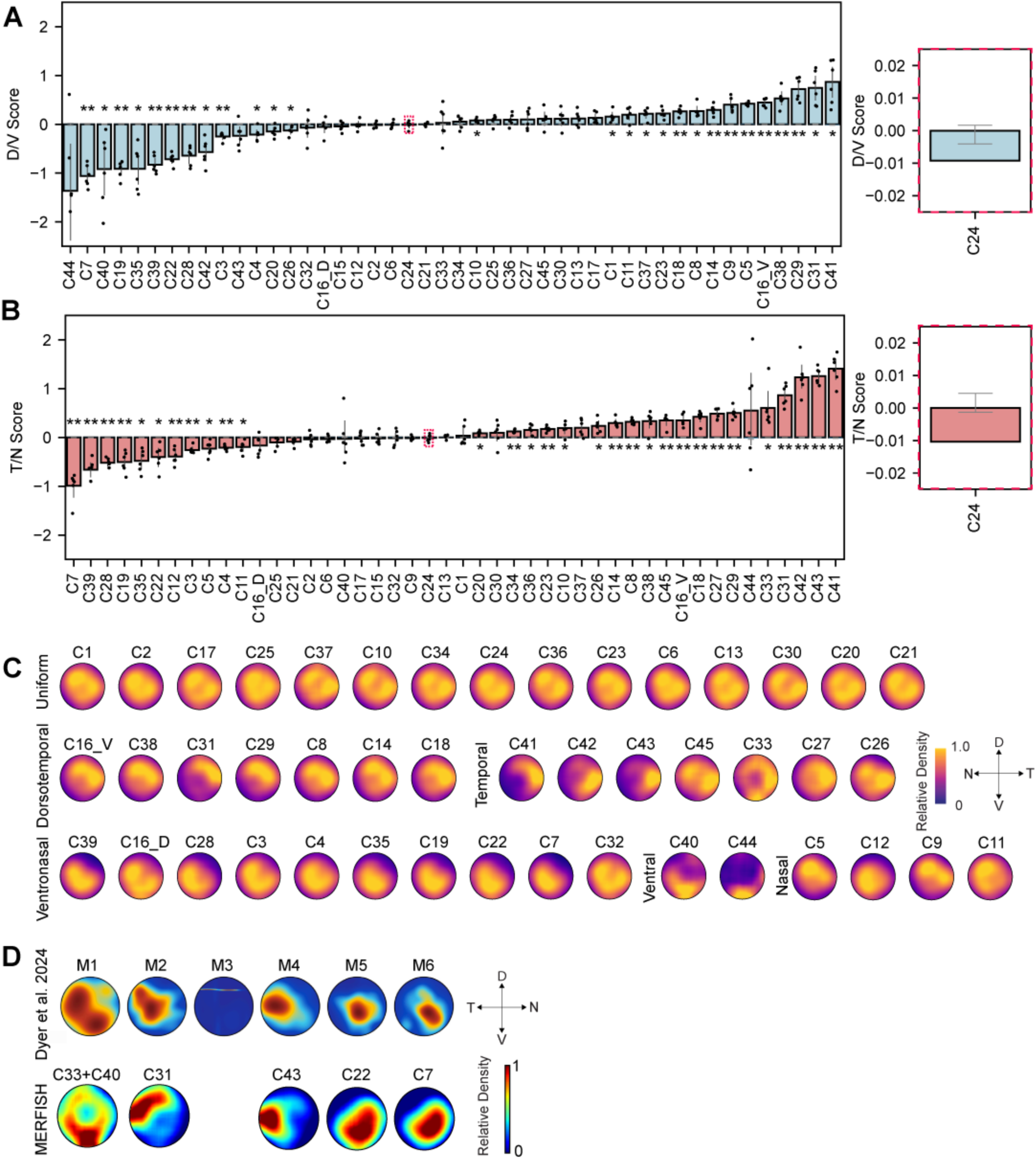
(Related to Figure 4) Topographic distributions for all 45 RGC types. A-B. D/V (A) and T/N (B) scores for each RGC type, calculated as *log2* changes in proportions across retinal halves. Black error bars, ± 1SD; grey error bars: null distribution SEM based on random permutations (e.g., C24 inset). The significance of rejecting the null hypothesis corresponding to a score of zero was assessed using a two-sided Welch’s t-test with the Benjamini-Hochberg correction (*p<0.05, **p<0.01). C. Comprehensive RGC topographic distributions, grouped by uniform, dorsal, ventral, temporal, and nasal biases. The adjusted density, which accounts for inhomogeneity in cell capture in an area, is plotted for a specific type (as in **Figure 4B**). D. Spatial distribution of putative ipRGC types to M1-M6 distributions, published in a recent study characterizing a newly established transgenic line in (Dyer et al., 2024a)

**Figure S6.**
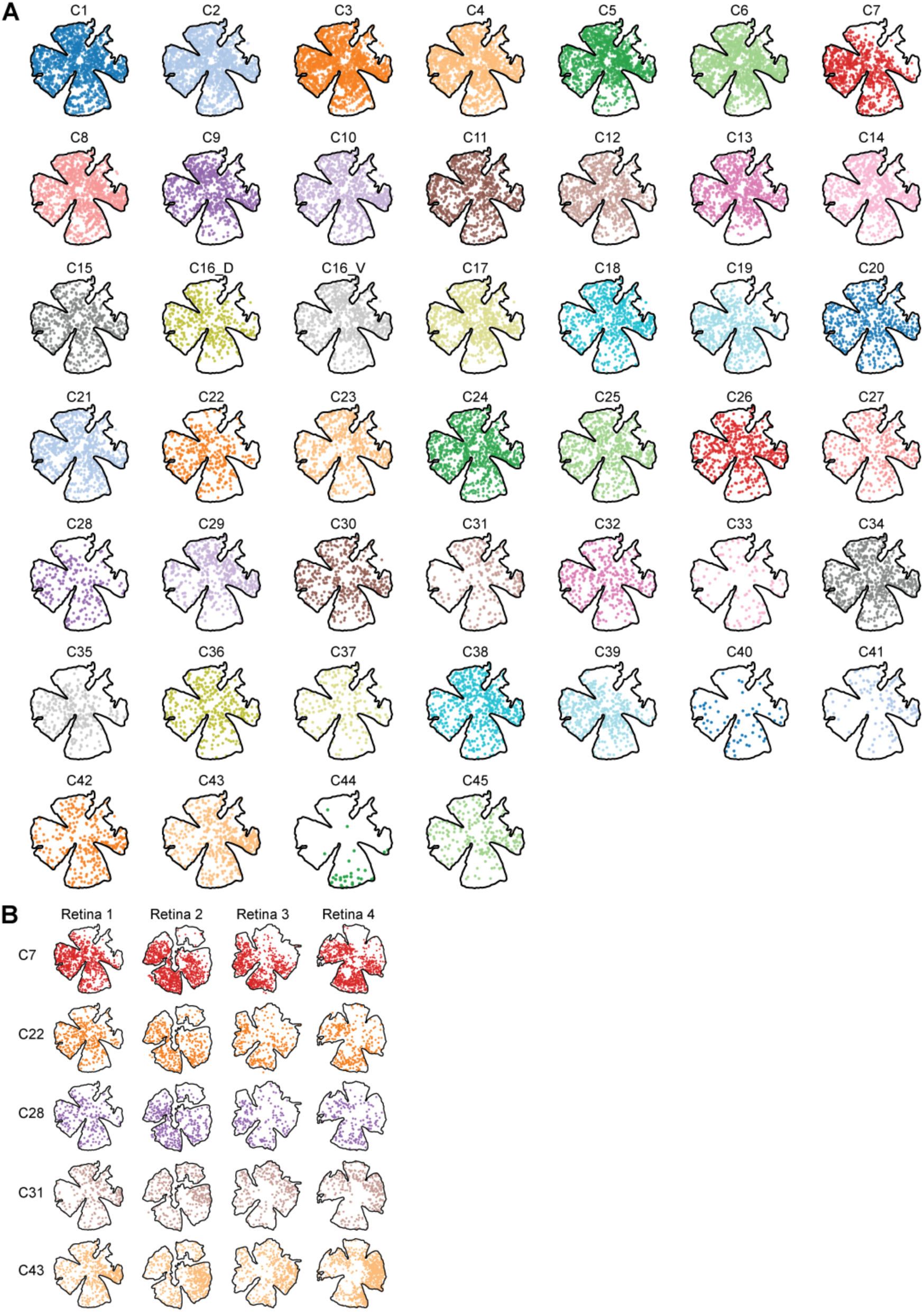
(Related to Figure 4) Raw data corresponding to topographies of 45 RGC types. A. Coordinates of all RGC types C1-C45 in a single retina. B. Coordinates of C7, C22, C28, C31, and C43 across four retinae, showing consistent topographic biases across biological replicates.

**Figure S7.**
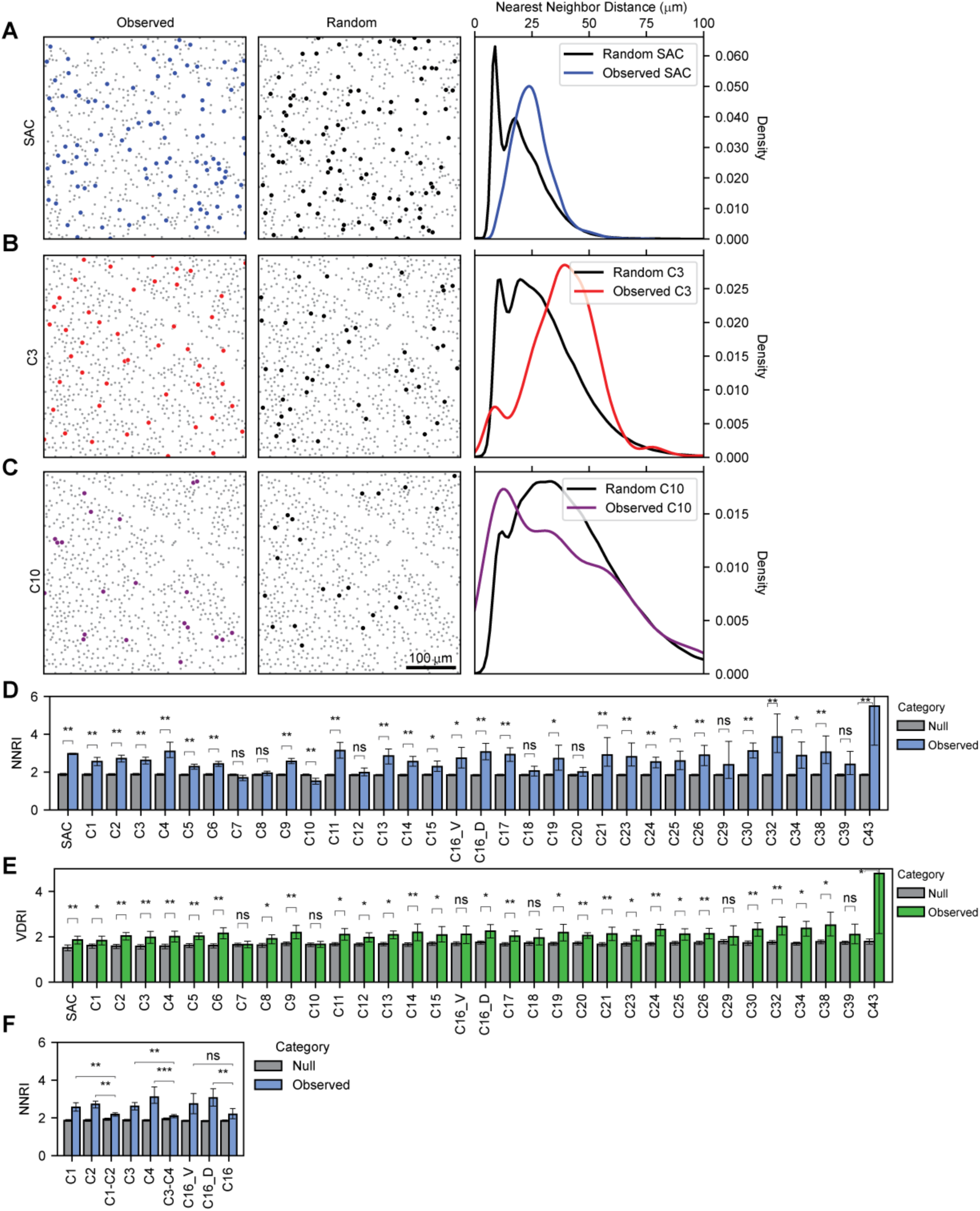
(Related to Figure 4) Mosaicism analyses to understand the regularity of 45 RGC types. A. Distribution of SACs: observed (left) and randomized (middle) for a select ROI. The right panel compares the observed vs. random distribution of nearest neighbor distances across multiple (n=9) ROIs. B. Same as A for C3 (F-RGCs), showing high regularity. C. Same as A for C10 (ON DSGCs), showing irregularity, as this contains 3 distinct (D, V, NT) types. Scale bars, 100 μm in A-C. D. Nearest Neighbor Regularity Index (NNRI) comparison for observed and null distributions of selected RGC types. Significance was assessed using the Wilcoxon rank-sum test with the Benjamini-Hochberg correction. (*p<0.05, **p<0.01) E. Voronoi Domain Regularity Index (VDRI) for RGC types, as in panel D. F. NNRI for individual cell types C1, C2, C3, C4, C16_D, and C16_V compared to artificial mixtures. Higher NNRI scores for individual types indicate stronger regularity (*p<0.05, **p<0.01, ***p<0.005).

**Figure S8.**
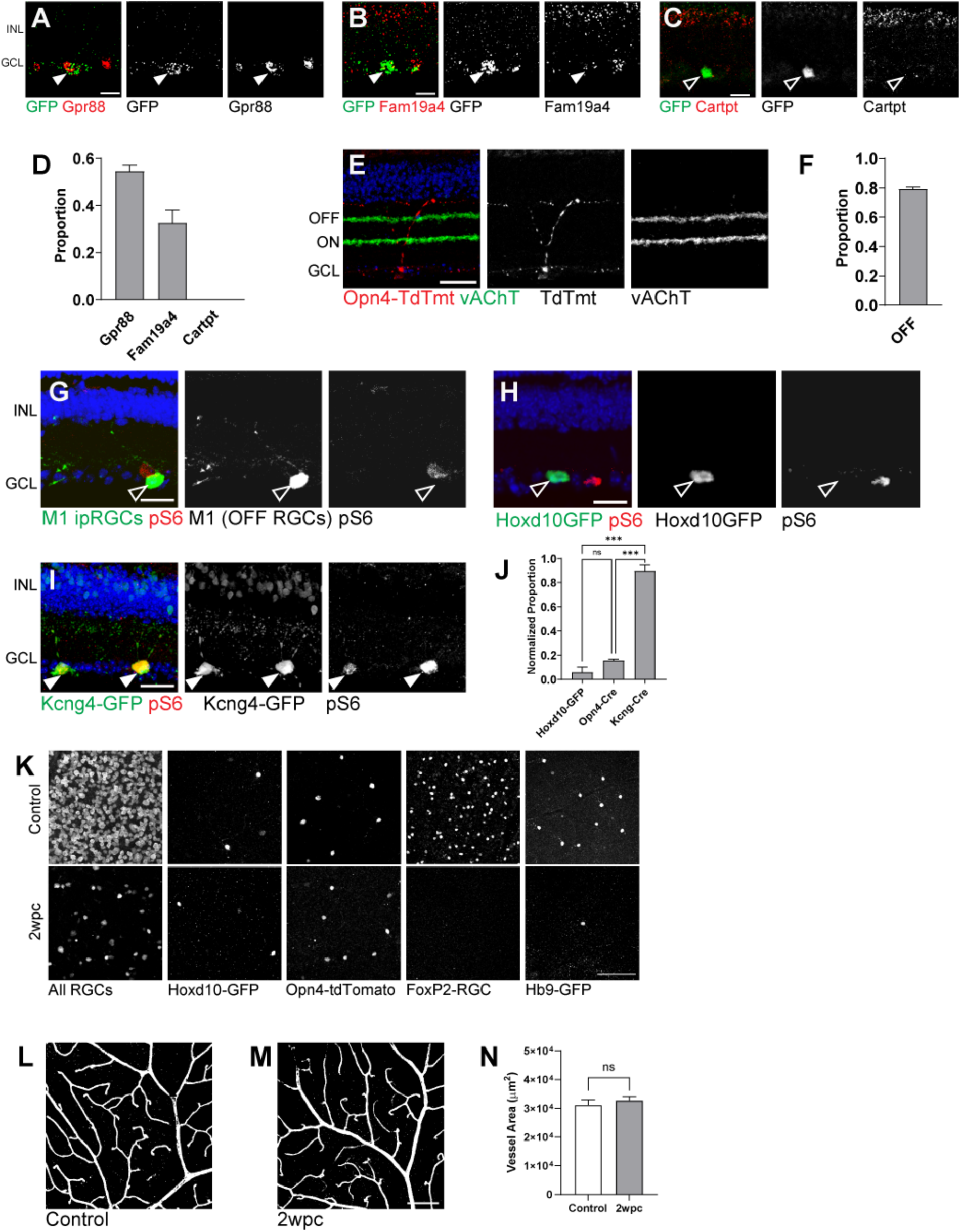
(Related to Figure 5) Immunohistochemical characterization of two perivascular RGC subclasses marking lines, including Opn4-TdTomato (ipRGCs) and Hoxd10-GFP (DSGCs). A-D. FISH and IHC characterizing Hox10-GFP RGCs including ON DSGCs (C10, A), as GFP^+^/Gpr88^+^ and GFP^+^/Fam19a4^+^(C24, B); but GFP^+^/Cartpt-negative (C16 and C12, C). Scale bars: 20µm. Arrowheads indicate double-positive cells; Open arrows indicate GFP+ only cells. Percentages of Hoxd10-GFP^+^ RGCs were quantified in (D), N= 4 animals in each condition. E-F. Opn4-TdTomato RGCs were characterized in the vertical retina section to be dominantly OFF-ipRGCs (M1-ipRGCs, C33 & C40) with dendrites projecting to the OFF-sublamina with a low fraction (of the rest) as ON-M2 ipRGCs. N = 4 animals in each condition. G-J. Sample images of Opn4-TdTomato RGCs (G), Hoxd10-GFP RGCs (H), and Kcng4-YFP RGCs (I) for pS6 staining to reflect mTOR activity. Quantification (J) of pS6 level was compared in three different RGC subclasses, indicating that DSGCs and ipRGCs possess lower mTOR activity than αRGCs. Arrowheads indicate pS6+ cells; Open arrows indicate pS6-negative cells. Scale bars: 20µm. N= 3 animals in each condition, One-Way ANOVA, p<0.0001 (****). K. Sample images of retinal flatmounts 2 weeks post ONC. Survival of Hoxd10-GFP (C10, C24) and Opn4-TdTomato (C40, C33) at 2 weeks post crush (wpc) was higher compared to all RGCs, Foxp2-RGCs(C3, C4, C28, C32, C38), and Hb9-GFP RGCs (C16), correlating with their perivascular distribution, as quantified in Figure 5O. Scale bars: 50µm. N = 4 animals in each condition. L-N. Sample images comparing the superficial layer (SL) of the retinal vasculature between control conditions (L) and 2wpc (M) conditions. Quantifications of the SL vascular densities (N) over an area of 0.28 mm^2^ indicated that ONC at 2wpc did not cause major vessel density changes. Scale bars: 100µm. Student’s T-test, N=4 retinas each, N.S., indicates no significance.

---

**Table S1.**
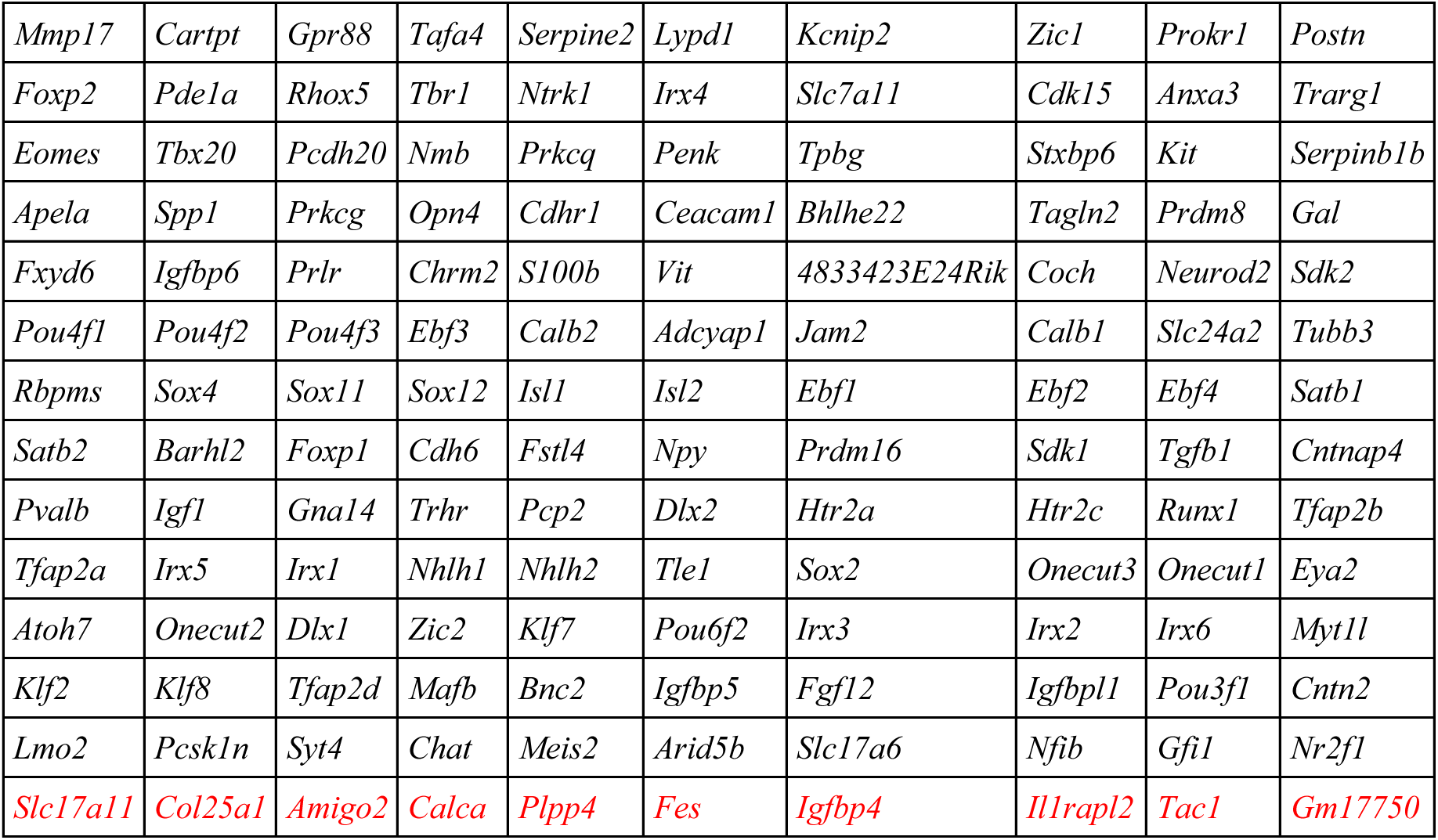
A pre-designed MERFISH 140-gene panel for mouse RGC type classification using MERSCOPE (Vizgen, VZG180) RGC type-specific marker genes were selected based on scRNA-Seq data. Among all 140 genes, we included several AC markers. In addition, ten marker genes in red (last row) were removed from the subsequent data analysis due to either low specificity of the transcript in marking RGC clusters or rare transcript detection in the adult RGC MERFISH datasets. The rest of the 130 marker genes were fully utilized in the current analysis.

**Table S2.**
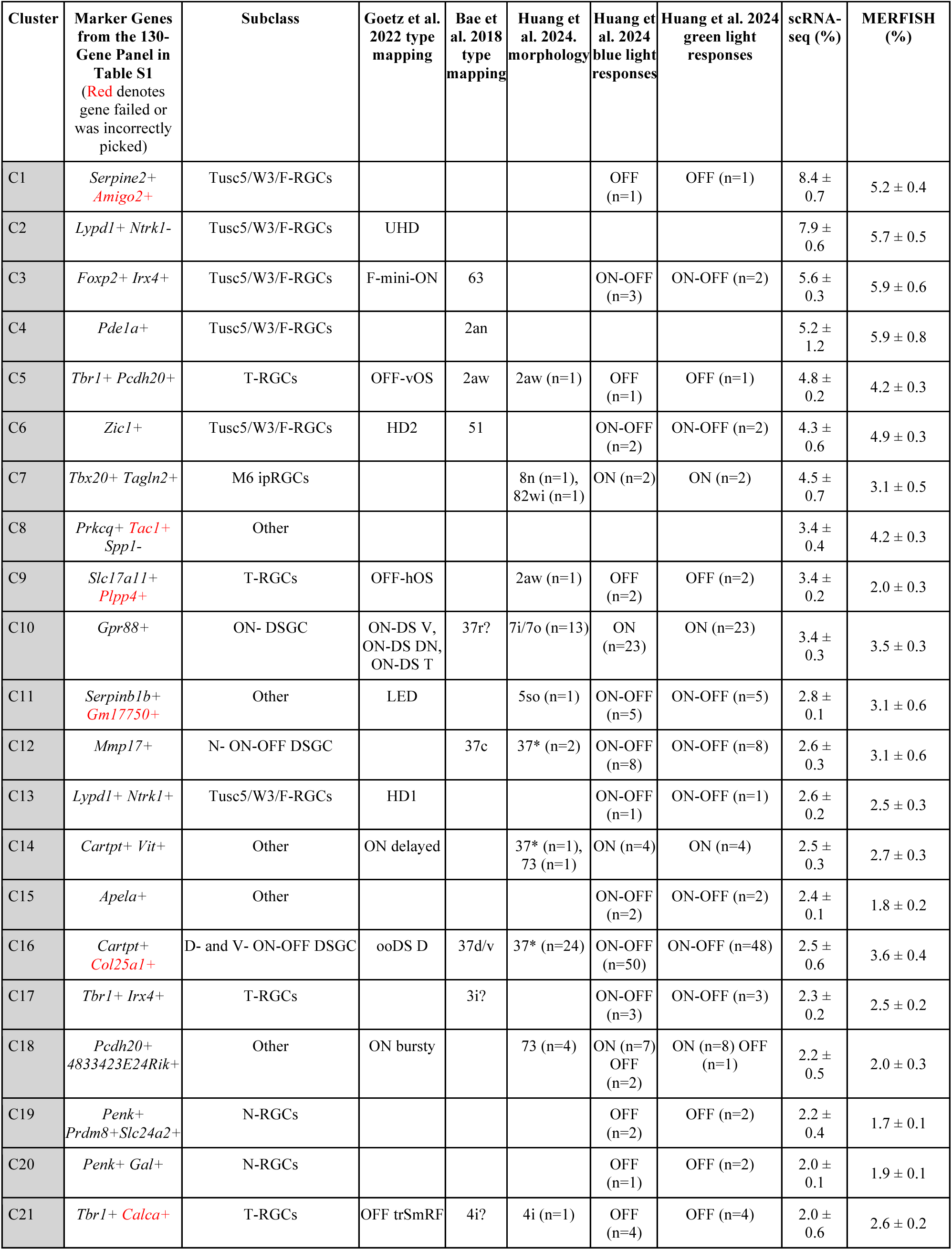
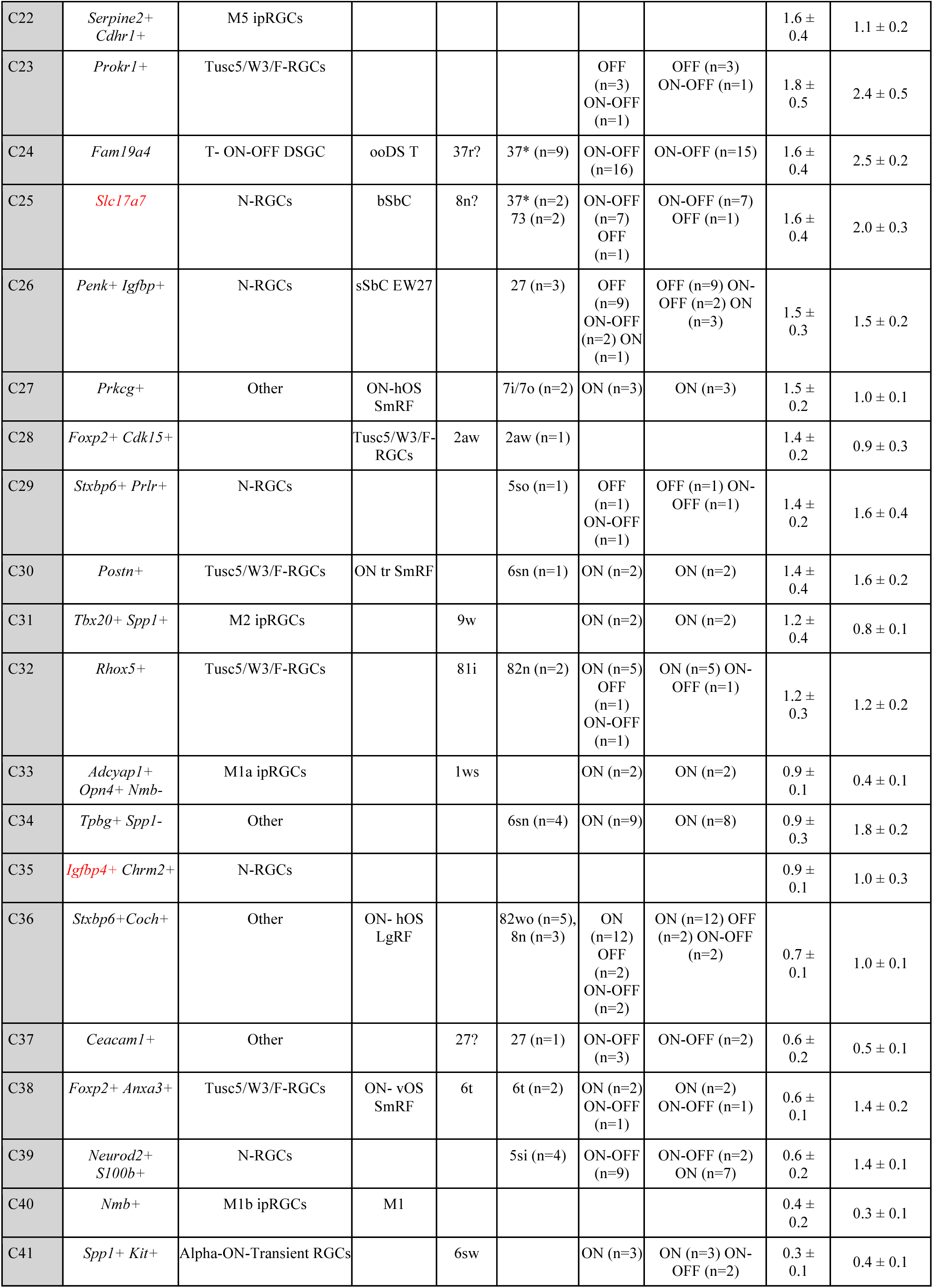
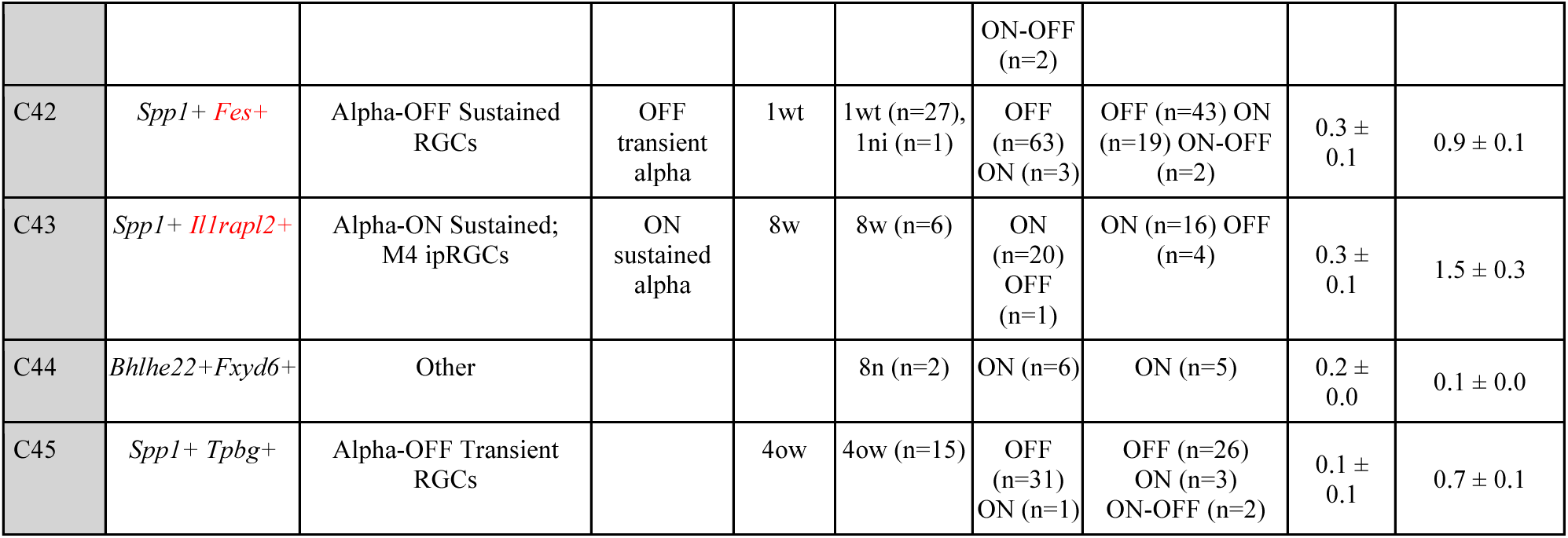
RGC Cluster frequencies comparison between scRNA-seq and MERFISH.

**Table S3.** Animal numbers, ages, and genotypes reported.

**Table S4.** All statistics and analysis methods reported.

