## Supplementary material for "Molecular and spatial analysis of ganglion cells on retinal flatmounts: diversity, topography, and perivascularity": Table S1

**Table S1. A pre-designed MERFISH 140-gene panel for mouse RGC type classification using MERSCOPE (Vizgen, VZG180)**

RGC type-specific marker genes were selected based on scRNA-Seq data. Among all 140 genes, we included several AC markers. In addition, multiple marker genes in red (last row) were removed from the subsequent data analysis due to either low specificity of the transcript in marking RGC clusters or rare transcript detection in the adult RGC MERFISH datasets. The rest of the 130 marker genes were fully utilized in the current analysis.

| | | | | | | | | | |
|---|---|---|---|---|---|---|---|---|---|
| *Mmp17* | *Cartpt* | *Gpr88* | *Tafa4* | *Serpine2* | *Lypd1* | *Kcnip2* | *Zic1* | *Prokr1* | *Postn* |
| *Foxp2* | *Pde1a* | *Rhox5* | *Tbr1* | *Ntrk1* | *Irx4* | *Slc7a11* | *Cdk15* | *Anxa3* | *Trarg1* |
| *Eomes* | *Tbx20* | *Pcdh20* | *Nmb* | *Prkcq* | *Penk* | *Tpbg* | *Stxbp6* | *Kit* | *Serpinb1b* |
| *Apela* | *Spp1* | *Prkcg* | *Opn4* | *Cdhr1* | *Ceacam1* | *Bhlhe22* | *Tagln2* | *Prdm8* | *Gal* |
| *Fxyd6* | *Igfbp6* | *Prlr* | *Chrm2* | *S100b* | *Vit* | *4833423E24Rik* | *Coch* | *Neurod2* | *Sdk2* |
| *Pou4f1* | *Pou4f2* | *Pou4f3* | *Ebf3* | *Calb2* | *Adcyap1* | *Jam2* | *Calb1* | *Slc24a2* | *Tubb3* |
| *Rbpms* | *Sox4* | *Sox11* | *Sox12* | *Isl1* | *Isl2* | *Ebf1* | *Ebf2* | *Ebf4* | *Satb1* |
| *Satb2* | *Barhl2* | *Foxp1* | *Cdh6* | *Fstl4* | *Npy* | *Prdm16* | *Sdk1* | *Tgfb1* | *Cntnap4* |
| *Pvalb* | *Igf1* | *Gna14* | *Trhr* | *Pcp2* | *Dlx2* | *Htr2a* | *Htr2c* | *Runx1* | *Tfap2b* |
| *Tfap2a* | *Irx5* | *Irx1* | *Nhlh1* | *Nhlh2* | *Tle1* | *Sox2* | *Onecut3* | *Onecut1* | *Eya2* |
| *Atoh7* | *Onecut2* | *Dlx1* | *Zic2* | *Klf7* | *Pou6f2* | *Irx3* | *Irx2* | *Irx6* | *Myt1l* |
| *Klf2* | *Klf8* | *Tfap2d* | *Mafb* | *Bnc2* | *Igfbp5* | *Fgf12* | *Igfbpl1* | *Pou3f1* | *Cntn2* |
| *Lmo2* | *Pcsk1n* | *Syt4* | *Chat* | *Meis2* | *Arid5b* | *Slc17a6* | *Nfib* | *Gfi1* | *Nr2f1* |
| *Slc17a11* | *Col25a1* | *Amigo2* | *Calca* | *Plpp4* | *Fes* | *Igfbp4* | *Il1rapl2* | *Tac1* | *Gm17750* |
