## Supplementary material for "Molecular and spatial analysis of ganglion cells on retinal flatmounts: diversity, topography, and perivascularity": Table S2

### Table S2. RGC Cluster comparison between scRNA-seq and MERFISH

| Cluster | Marker Genes from the 130-Gene Panel in Table S1 (Red denotes gene failed or was incorrectly picked) | Subclass | Goetz et al. 2022 type mapping | Bae et al. 2018 type mapping | Huang et al. 2024. morphology | Huang et al. 2024 blue light responses | Huang et al. 2024 green light responses | scRNA-seq (%) | MERFISH (%) |
|---|---|---|---|---|---|---|---|---|---|
| C1 | *Serpine2+ Amigo2+* | Tusc5/W3/F-RGCs | | | | OFF (n=1) | OFF (n=1) | 8.4 ± 0.7 | 5.2 ± 0.4 |
| C2 | *Lypd1+ Ntrk1-* | Tusc5/W3/F-RGCs | UHD | | | | | 7.9 ± 0.6 | 5.7 ± 0.5 |
| C3 | *Foxp2+ Irx4+* | Tusc5/W3/F-RGCs | F-mini-ON | 63 | | ON-OFF (n=3) | ON-OFF (n=2) | 5.6 ± 0.3 | 5.9 ± 0.6 |
| C4 | *Pde1a+* | Tusc5/W3/F-RGCs | | 2an | | | | 5.2 ± 1.2 | 5.9 ± 0.8 |
| C5 | *Tbr1+ Pcdh20+* | T-RGCs | OFF-vOS | 2aw | 2aw (n=1) | OFF (n=1) | OFF (n=1) | 4.8 ± 0.2 | 4.2 ± 0.3 |
| C6 | *Zic1+* | Tusc5/W3/F-RGCs | HD2 | 51 | | ON-OFF (n=2) | ON-OFF (n=2) | 4.3 ± 0.6 | 4.9 ± 0.3 |
| C7 | *Tbx20+ Tagln2+* | M6 ipRGCs | | | 8n (n=1), 82wi (n=1) | ON (n=2) | ON (n=2) | 4.5 ± 0.7 | 3.1 ± 0.5 |
| C8 | *Prkcq+ Tac1+ Spp1-* | Other | | | | | | 3.4 ± 0.4 | 4.2 ± 0.3 |
| C9 | *Slc17a11+ Plpp4+* | T-RGCs | OFF-hOS | | 2aw (n=1) | OFF (n=2) | OFF (n=2) | 3.4 ± 0.2 | 2.0 ± 0.3 |
| C10 | *Gpr88+* | ON- DSGC | ON-DS V, ON-DS DN, ON-DS T | 37r? | 7i/7o (n=13) | ON (n=23) | ON (n=23) | 3.4 ± 0.3 | 3.5 ± 0.3 |
| C11 | *Serpinb1b+ Gm17750+* | Other | LED | | 5so (n=1) | ON-OFF (n=5) | ON-OFF (n=5) | 2.8 ± 0.1 | 3.1 ± 0.6 |
| C12 | *Mmp17+* | N- ON-OFF DSGC | | 37c | 37* (n=2) | ON-OFF (n=8) | ON-OFF (n=8) | 2.6 ± 0.3 | 3.1 ± 0.6 |
| C13 | *Lypd1+ Ntrk1+* | Tusc5/W3/F-RGCs | HD1 | | | ON-OFF (n=1) | ON-OFF (n=1) | 2.6 ± 0.2 | 2.5 ± 0.3 |
| C14 | *Cartpt+ Vit+* | Other | ON delayed | | 37* (n=1), 73 (n=1) | ON (n=4) | ON (n=4) | 2.5 ± 0.3 | 2.7 ± 0.3 |
| C15 | *Apela+* | Other | | | | ON-OFF (n=2) | ON-OFF (n=2) | 2.4 ± 0.1 | 1.8 ± 0.2 |
| C16 | *Cartpt+ Col25a1+* | D- and V- ON-OFF DSGC | ooDS D | 37d/v | 37* (n=24) | ON-OFF (n=50) | ON-OFF (n=48) | 2.5 ± 0.6 | 3.6 ± 0.4 |
| C17 | *Tbr1+ Irx4+* | T-RGCs | | 3i? | | ON-OFF (n=3) | ON-OFF (n=3) | 2.3 ± 0.2 | 2.5 ± 0.2 |
| C18 | *Pcdh20+ 4833423E24Rik+* | Other | ON bursty | | 73 (n=4) | ON (n=7) OFF (n=2) | ON (n=8) OFF (n=1) | 2.2 ± 0.5 | 2.0 ± 0.3 |
| C19 | *Penk+ Prdm8+Slc24a2+* | N-RGCs | | | | OFF (n=2) | OFF (n=2) | 2.2 ± 0.4 | 1.7 ± 0.1 |
| C20 | *Penk+ Gal+* | N-RGCs | | | | OFF (n=1) | OFF (n=2) | 2.0 ± 0.1 | 1.9 ± 0.1 |
| C21 | *Tbr1+ Calca+* | T-RGCs | OFF trSmRF | 4i? | 4i (n=1) | OFF (n=4) | OFF (n=4) | 2.0 ± 0.6 | 2.6 ± 0.2 |

| | | | | | | | | | |
|---|---|---|---|---|---|---|---|---|---|
| C22 | *Serpine2+ Cdhr1+* | M5 ipRGCs | | | | | | 1.6 ± 0.4 | 1.1 ± 0.2 |
| C23 | *Prokr1+* | Tusc5/W3/F-RGCs | | | | OFF (n=3) ON-OFF (n=1) | OFF (n=3) ON-OFF (n=1) | 1.8 ± 0.5 | 2.4 ± 0.5 |
| C24 | *Fam19a4* | T- ON-OFF DSGC | ooDS T | 37r? | 37* (n=9) | ON-OFF (n=16) | ON-OFF (n=15) | 1.6 ± 0.4 | 2.5 ± 0.2 |
| C25 | *Slc17a7* | N-RGCs | bSbC | 8n? | 37* (n=2) 73 (n=2) | ON-OFF (n=7) OFF (n=1) | ON-OFF (n=7) OFF (n=1) | 1.6 ± 0.4 | 2.0 ± 0.3 |
| C26 | *Penk+ Igfbp+* | N-RGCs | sSbC EW27 | | 27 (n=3) | OFF (n=9) ON-OFF (n=2) ON (n=1) | OFF (n=9) ON-OFF (n=2) ON (n=3) | 1.5 ± 0.3 | 1.5 ± 0.2 |
| C27 | *Prkcg+* | Other | ON-hOS SmRF | | 7i/7o (n=2) | ON (n=3) | ON (n=3) | 1.5 ± 0.2 | 1.0 ± 0.1 |
| C28 | *Foxp2+ Cdk15+* | | Tusc5/W3/F-RGCs | 2aw | 2aw (n=1) | | | 1.4 ± 0.2 | 0.9 ± 0.3 |
| C29 | *Stxbp6+ Prlr+* | N-RGCs | | | 5so (n=1) | OFF (n=1) ON-OFF (n=1) | OFF (n=1) ON-OFF (n=1) | 1.4 ± 0.2 | 1.6 ± 0.4 |
| C30 | *Postn+* | Tusc5/W3/F-RGCs | ON tr SmRF | | 6sn (n=1) | ON (n=2) | ON (n=2) | 1.4 ± 0.4 | 1.6 ± 0.2 |
| C31 | *Tbx20+ Spp1+* | M2 ipRGCs | | 9w | | ON (n=2) | ON (n=2) | 1.2 ± 0.4 | 0.8 ± 0.1 |
| C32 | *Rhox5+* | Tusc5/W3/F-RGCs | | 81i | 82n (n=2) | ON (n=5) OFF (n=1) ON-OFF (n=1) | ON (n=5) ON-OFF (n=1) | 1.2 ± 0.3 | 1.2 ± 0.2 |
| C33 | *Adcyap1+ Opn4+ Nmb-* | M1a ipRGCs | | 1ws | | ON (n=2) | ON (n=2) | 0.9 ± 0.1 | 0.4 ± 0.1 |
| C34 | *Tpbg+ Spp1-* | Other | | | 6sn (n=4) | ON (n=9) | ON (n=8) | 0.9 ± 0.3 | 1.8 ± 0.2 |
| C35 | *Igfbp4+ Chrm2+* | N-RGCs | | | | | | 0.9 ± 0.1 | 1.0 ± 0.3 |
| C36 | *Stxbp6+Coch+* | Other | ON- hOS LgRF | | 82wo (n=5), 8n (n=3) | ON (n=12) OFF (n=2) ON-OFF (n=2) | ON (n=12) OFF (n=2) ON-OFF (n=2) | 0.7 ± 0.1 | 1.0 ± 0.1 |
| C37 | *Ceacam1+* | Other | | 27? | 27 (n=1) | ON-OFF (n=3) | ON-OFF (n=2) | 0.6 ± 0.2 | 0.5 ± 0.1 |
| C38 | *Foxp2+ Anxa3+* | Tusc5/W3/F-RGCs | ON- vOS SmRF | 6t | 6t (n=2) | ON (n=2) ON-OFF (n=1) | ON (n=2) ON-OFF (n=1) | 0.6 ± 0.1 | 1.4 ± 0.2 |
| C39 | *Neurod2+ S100b+* | N-RGCs | | | 5si (n=4) | ON-OFF (n=9) | ON-OFF (n=2) ON (n=7) | 0.6 ± 0.2 | 1.4 ± 0.1 |
| C40 | *Nmb+* | M1b ipRGCs | M1 | | | | | 0.4 ± 0.2 | 0.3 ± 0.1 |
| C41 | *Spp1+ Kit+* | Alpha-ON-Transient RGCs | | 6sw | | ON (n=3) ON-OFF (n=2) | ON (n=3) ON-OFF (n=2) | 0.3 ± 0.1 | 0.4 ± 0.1 |
| C42 | *Spp1+ Fes+* | Alpha-OFF Sustained RGCs | OFF transient alpha | 1wt | 1wt (n=27), 1ni (n=1) | OFF (n=63) ON (n=3) | OFF (n=43) ON (n=19) ON-OFF (n=2) | 0.3 ± 0.1 | 0.9 ± 0.1 |

| | | | | | | | | | |
|---|---|---|---|---|---|---|---|---|---|
| C43 | *Spp1+ Il1rapl2+* | Alpha-ON Sustained; M4 ipRGCs | ON sustained alpha | 8w | 8w (n=6) | ON (n=20) OFF (n=1) | ON (n=16) OFF (n=4) | 0.3 ± 0.1 | 1.5 ± 0.3 |
| C44 | *Bhlhe22+Fxyd6+* | Other | | | 8n (n=2) | ON (n=6) | ON (n=5) | 0.2 ± 0.0 | 0.1 ± 0.0 |
| C45 | *Spp1+ Tpbg+* | Alpha-OFF Transient RGCs | | 4ow | 4ow (n=15) | OFF (n=31) ON (n=1) | OFF (n=26) ON (n=3) ON-OFF (n=2) | 0.1 ± 0.1 | 0.7 ± 0.1 |
