## Supplementary material for "Molecular and spatial analysis of ganglion cells on retinal flatmounts: diversity, topography, and perivascularity": Table S3

**Table S3  Summary of all experimental designs**

| Figures | Aims | Mouse lines | Manipulations | Age | Type of data |
|---|---|---|---|---|---|
| Figure 1E-H | Preparation of retinal flatmounts | C57BL/6 (Wildtype) | N/A | P100 | MERFISH |
| Figure 2A-F | Characterization of cell types in GCL | | N/A | P100 | MERFISH |
| Figure 3A-E | Profiling of RGC types | | N/A | P78 | MERFISH |
| Figure 4A-B | Identification of RGC type gradients through MERFISH | | N/A | P78 | MERFISH |
| | | | N/A | P115 | MERFISH |
| | | | N/A | P115 | MERFISH |
| | | | | 6 Replicates | |
| Figures S1,S2,S3,S5,S6,S7 | Additional analyses of MERFISH data | ` | N/A | | |
| Figure S4, A-D | Merfish characterization of Melanopsin+ RGCs | | N/A | P135 | MERFISH |
| Figure 2G-I | MERFISH cell type correspondence with CD31 staining | C57BL/6 (Wildtype) | N/A | P100 | MERFISH |
| Figure 3F-K | Dendrite reconstruction of Melanopsin+ RGCs | Opn4-Cre;LSL-YFP | N/A | P184 | IHC |
| Figure 4C-F | Opn4 & Nmb Flatmount ISH | | N/A | P72-108 | ISH |
| Figure 5C | Identification of perivascular RGC types | C57BL/6 (Wildtype) | N/A | | MERFISH |
| Figure 5D-H | Shortest distance analysis of RGC types to vessels | Hoxd10-GFP | N/A | P30-P70 | IHC |
| | | Nts-Cre; LSL-GFP | N/A | P65 | IHC |
| | | Opn4-tdTomato | N/A | P60-P80 | IHC |
| | | Hb9-GFP | N/A | P60-P145 | IHC |
| Figure 5J-M | Perivascular RGC types in human | N/A | N/A | GW22-23 | IHC |
| Figure 5O | Survival of RGC types 2 weeks post optic nerve crush | Same Figure S8K | ONC 2WPC | | |
| Figure S8K | Vessel area quantification in control and 2 weeks post optic nerve crush | Opn4-tdTomato | ONC 2WPC | P120 | IHC |
| | | Hb9-GFP | ONC 2WPC | P224 | IHC |
| | | Hoxd10-GFP | ONC 2WPC | P120 | IHC |
| Figure S8, L, M, N | | C57BL/6 (Wildtype) | ONC 2WPC | P86 | IHC |
| Figure S8, A-D | Characterization of Hoxd10-GFP line | Hoxd10-GFP | N/A | P20-P59 | ISH |
| Figure S8, E-F | Characterization of Opn4-tdTomato line | Opn4-tdTomato | N/A | P60-P80 | IHC |
| Figure S8,G, J | PS6 staining of RGC reporter lines | Opn4-tdTomato | N/A | P60-P80 | IHC |
| Figure S8,H, J | | Hoxd10-GFP | N/A | P20-P59 | IHC |
| Figure S8,I, J | | Kcng4-Cre;LSL-YFP | N/A | P133 | IHC |
