## Supplementary material for "Molecular and spatial analysis of ganglion cells on retinal flatmounts: diversity, topography, and perivascularity": Table S4

### Table S4 Summary of statistical analyses

| Figure | Sample Size | Statistical test | Result (p value) | |
|---|---|---|---|---|
| Figure 4A | 6 | Two-sided Welch's t-test + Benjamini Hochberg Correction | D/V Score | |
| | | | C1 (Observed vs. Random) | 0.033 |
| | | | C2 (Observed vs. Random) | 0.624 |
| | | | C3 (Observed vs. Random) | 0.003 |
| | | | C4 (Observed vs. Random) | 0.022 |
| | | | C5 (Observed vs. Random) | 0.000 |
| | | | C6 (Observed vs. Random) | 0.806 |
| | | | C7 (Observed vs. Random) | 0.001 |
| | | | C8 (Observed vs. Random) | 0.009 |
| | | | C9 (Observed vs. Random) | 0.003 |
| | | | C10 (Observed vs. Random) | 0.011 |
| | | | C11 (Observed vs. Random) | 0.031 |
| | | | C12 (Observed vs. Random) | 0.775 |
| | | | C13 (Observed vs. Random) | 0.156 |
| | | | C14 (Observed vs. Random) | 0.003 |
| | | | C15 (Observed vs. Random) | 0.374 |
| | | | C16_D (Observed vs. Random) | 0.526 |
| | | | C16_V (Observed vs. Random) | 0.001 |
| | | | C17 (Observed vs. Random) | 0.053 |
| | | | C18 (Observed vs. Random) | 0.003 |
| | | | C19 (Observed vs. Random) | 0.001 |
| | | | C20 (Observed vs. Random) | 0.039 |
| | | | C21 (Observed vs. Random) | 0.943 |
| | | | C22 (Observed vs. Random) | 0.001 |
| | | | C23 (Observed vs. Random) | 0.008 |
| | | | C24 (Observed vs. Random) | 0.807 |
| | | | C25 (Observed vs. Random) | 0.114 |
| | | | C26 (Observed vs. Random) | 0.027 |
| | | | C27 (Observed vs. Random) | 0.338 |
| | | | C28 (Observed vs. Random) | 0.003 |
| | | | C29 (Observed vs. Random) | 0.003 |
| | | | C30 (Observed vs. Random) | 0.253 |
| | | | C31 (Observed vs. Random) | 0.009 |
| | | | C32 (Observed vs. Random) | 0.624 |
| | | | C33 (Observed vs. Random) | 0.858 |
| | | | C34 (Observed vs. Random) | 0.403 |
| | | | C35 (Observed vs. Random) | 0.010 |
| | | | C36 (Observed vs. Random) | 0.159 |
| | | | C37 (Observed vs. Random) | 0.010 |
| | | | C38 (Observed vs. Random) | 0.003 |
| | | | C39 (Observed vs. Random) | 0.001 |
| | | | C40 (Observed vs. Random) | 0.048 |
| | | | C41 (Observed vs. Random) | 0.011 |
| | | | C42 (Observed vs. Random) | 0.009 |
| | | | C43 (Observed vs. Random) | 0.109 |
| | | | C44 (Observed vs. Random) | 0.085 |
| | | | C45 (Observed vs. Random) | 0.247 |
| | | | T/N Score | |
| | | | C1 (Observed vs. Random) | 0.781 |
| | | | C2 (Observed vs. Random) | 0.562 |
| | | | C3 (Observed vs. Random) | 0.001 |
| | | | C4 (Observed vs. Random) | 0.001 |
| | | | C5 (Observed vs. Random) | 0.020 |
| | | | C6 (Observed vs. Random) | 0.421 |
| | | | C7 (Observed vs. Random) | 0.001 |
| | | | C8 (Observed vs. Random) | 0.001 |
| | | | C9 (Observed vs. Random) | 0.562 |
| | | | C10 (Observed vs. Random) | 0.026 |
| | | | C11 (Observed vs. Random) | 0.019 |
| | | | C12 (Observed vs. Random) | 0.005 |

| | | | | |
|---|---|---|---|---|
| | | | C13 (Observed vs. Random) | 0.954 |
| | | | C14 (Observed vs. Random) | 0.001 |
| | | | C15 (Observed vs. Random) | 0.781 |
| | | | C16_D (Observed vs. Random) | 0.139 |
| | | | C16_V (Observed vs. Random) | 0.002 |
| | | | C17 (Observed vs. Random) | 0.781 |
| | | | C18 (Observed vs. Random) | 0.001 |
| | | | C19 (Observed vs. Random) | 0.005 |
| | | | C20 (Observed vs. Random) | 0.011 |
| | | | C21 (Observed vs. Random) | 0.060 |
| | | | C22 (Observed vs. Random) | 0.019 |
| | | | C23 (Observed vs. Random) | 0.002 |
| | | | C24 (Observed vs. Random) | 0.784 |
| | | | C25 (Observed vs. Random) | 0.099 |
| | | | C26 (Observed vs. Random) | 0.015 |
| | | | C27 (Observed vs. Random) | 0.001 |
| | | | C28 (Observed vs. Random) | 0.001 |
| | | | C29 (Observed vs. Random) | 0.001 |
| | | | C30 (Observed vs. Random) | 0.464 |
| | | | C31 (Observed vs. Random) | 0.001 |
| | | | C32 (Observed vs. Random) | 0.875 |
| | | | C33 (Observed vs. Random) | 0.025 |
| | | | C34 (Observed vs. Random) | 0.001 |
| | | | C35 (Observed vs. Random) | 0.008 |
| | | | C36 (Observed vs. Random) | 0.026 |
| | | | C37 (Observed vs. Random) | 0.052 |
| | | | C38 (Observed vs. Random) | 0.016 |
| | | | C39 (Observed vs. Random) | 0.002 |
| | | | C40 (Observed vs. Random) | 0.902 |
| | | | C41 (Observed vs. Random) | 0.001 |
| | | | C42 (Observed vs. Random) | 0.001 |
| | | | C43 (Observed vs. Random) | 0.000 |
| | | | C44 (Observed vs. Random) | 0.304 |
| | | | C45 (Observed vs. Random) | 0.002 |
| Figure 5C | 6 | Permutation test + Benjamini Hochberg correction | Edges between cell types (observed vs. null) | |
| | | | C10 to Non-neuronal | 2.00E-05 |
| | | | C8 to Non-neuronal | 0.0005 |
| | | | C24 to Non-neuronal | 0.0019 |
| | | | C43 to Non-neuronal | 0.002 |
| | | | C41 to Non-neuronal | 0.002 |
| | | | C33 to Non-neuronal | 0.006 |
| | | | C40 to Non-neuronal | 0.006 |
| | | | C31 to Non-neuronal | 0.014 |
| | | | C21 to Non-neuronal | 0.026 |
| | | | C18 to Non-neuronal | 0.11 |
| | | | C44 to Non-neuronal | 0.14 |
| | | | C29 to Non-neuronal | 0.21 |
| | | | C23 to Non-neuronal | 0.22 |
| | | | C16 to Non-neuronal | 0.26 |
| | | | C27 to Non-neuronal | 0.37 |
| | | | C34 to Non-neuronal | 0.58 |
| | | | C22 to Non-neuronal | 0.7 |
| | | | C30 to Non-neuronal | 0.74 |
| | | | C42 to Non-neuronal | 0.84 |
| | | | C45 to Non-neuronal | 0.84 |
| | | | C37 to Non-neuronal | 0.89 |
| | | | C38 to Non-neuronal | 0.89 |
| | | | C20 to Non-neuronal | 0.89 |
| | | | C25 to Non-neuronal | 0.95 |
| | | | C5 to Non-neuronal | >0.99 |
| | | | C39 to Non-neuronal | >0.99 |

| | | | | |
|---|---|---|---|---|
| | | | C6 to Non-neuronal | >0.99 |
| | | | C7 to Non-neuronal | >0.99 |
| | | | C4 to Non-neuronal | >0.99 |
| | | | C1 to Non-neuronal | >0.99 |
| | | | C3 to Non-neuronal | >0.99 |
| | | | C35 to Non-neuronal | >0.99 |
| | | | C32 to Non-neuronal | >0.99 |
| | | | C28 to Non-neuronal | >0.99 |
| | | | C26 to Non-neuronal | >0.99 |
| | | | C2 to Non-neuronal | >0.99 |
| | | | C19 to Non-neuronal | >0.99 |
| | | | C17 to Non-neuronal | >0.99 |
| | | | C15 to Non-neuronal | >0.99 |
| | | | C14 to Non-neuronal | >0.99 |
| | | | C13 to Non-neuronal | >0.99 |
| | | | C12 to Non-neuronal | >0.99 |
| | | | C11 to Non-neuronal | >0.99 |
| | | | C36 to Non-neuronal | >0.99 |
| | | | C9 to Non-neuronal | >0.99 |
| Figure 5D-H | 4 | Two-Way ANOVA | 0 to 1 µm | |
| | | | Hoxd10-GFP vs. Nts-GFP | 0.6657 |
| | | | Hoxd10-GFP vs. Opn4-tdTomato | 0.6036 |
| | | | Hoxd10-GFP vs. Foxp2-IHC | <0.0001 |
| | | | Hoxd10-GFP vs. Hb9-GFP | <0.0001 |
| | | | Nts-GFP vs. Opn4-tdTomato | >0.9999 |
| | | | Nts-GFP vs. Foxp2-IHC | <0.0001 |
| | | | Nts-GFP vs. Hb9-GFP | 0.0059 |
| | | | Opn4-tdTomato vs. Foxp2-IHC | <0.0001 |
| | | | Opn4-tdTomato vs. Hb9-GFP | 0.0025 |
| | | | Foxp2-IHC vs. Hb9-GFP | 0.3357 |
| | | | 1 to 10 µm | |
| | | | Hoxd10-GFP vs. Nts-GFP | >0.9999 |
| | | | Hoxd10-GFP vs. Opn4-tdTomato | 0.9979 |
| | | | Hoxd10-GFP vs. Foxp2-IHC | 0.0004 |
| | | | Hoxd10-GFP vs. Hb9-GFP | 0.0003 |
| | | | Nts-GFP vs. Opn4-tdTomato | 0.9986 |
| | | | Nts-GFP vs. Foxp2-IHC | 0.0011 |
| | | | Nts-GFP vs. Hb9-GFP | 0.0009 |
| | | | Opn4-tdTomato vs. Foxp2-IHC | 0.001 |
| | | | Opn4-tdTomato vs. Hb9-GFP | 0.0008 |
| | | | Foxp2-IHC vs. Hb9-GFP | >0.9999 |
| | | | Greater than 10 µm | |
| | | | Hoxd10-GFP vs. Nts-GFP | 0.9915 |
| | | | Hoxd10-GFP vs. Opn4-tdTomato | 0.7878 |
| | | | Hoxd10-GFP vs. Foxp2-IHC | 0.0655 |
| | | | Hoxd10-GFP vs. Hb9-GFP | 0.9428 |
| | | | Nts-GFP vs. Opn4-tdTomato | 0.9738 |
| | | | Nts-GFP vs. Foxp2-IHC | 0.2454 |
| | | | Nts-GFP vs. Hb9-GFP | 0.9991 |
| | | | Opn4-tdTomato vs. Foxp2-IHC | 0.5051 |
| | | | Opn4-tdTomato vs. Hb9-GFP | 0.9953 |
| | | | Foxp2-IHC vs. Hb9-GFP | 0.2918 |
| Figure 5J-M | 3 | One-Way ANOVA | Opn4 vs. Bnc2 | 0.6343 |
| | | | Opn4 vs. Foxp2 | <0.0001 |
| | | | Bnc2 vs. Foxp2 | <0.0001 |
| Figure 5O | 4 | One-Way ANOVA | Hoxd10-GFP vs. Opn4-tdTomato | 0.0022 |

| | | | |
|---|---|---|---|
| | | Hoxd10-GFP vs. Hb9-GFP | <0.0001 |
| | | Hoxd10-GFP vs. Foxp2-IHC | <0.0001 |
| | | Opn4-tdTomato vs. Hb9-GFP | <0.0001 |
| | | Opn4-tdTomato vs. Foxp2-IHC | <0.0001 |
| | | Hb9-GFP vs. Foxp2-IHC | 0.0905 |
| | | Hb9-GFP vs. All RGCs | <0.0001 |
| Figure S8K, 4 | Student's t-test | 2wpc vs. Control | 0.51 |
| Figure S8G, 3 | One-Way ANOVA | Hoxd10-GFP vs. Opn4-Cre | 0.2756 |
| | | Hoxd10-GFP vs. Kcng-Cre | <0.0001 |
| | | Opn4-Cre vs. Kcng-Cre | <0.0001 |
